# Nephronophthisis-associated FBW7 mediates cyst-dependent decline of renal function in ADPKD

**DOI:** 10.1101/2024.02.29.582788

**Authors:** Maulin Mukeshchandra Patel, Vasileios Gerakopoulos, Eleni Petsouki, Kurt A. Zimmerman, Leonidas Tsiokas

## Abstract

Nephronophthisis (NPHP) and autosomal dominant Polycystic Kidney Disease (ADPKD) are two genetically distinct forms of Polycystic Kidney Disease (PKD), yet both diseases present with kidney cysts and a gradual decline in renal function. Prevailing dogma in PKD is that changes in kidney architecture account for the decline in kidney function, but the molecular/cellular basis of such coupling is unknown. To address this question, we induced a form of proteome reprogramming by deleting *Fbxw7* encoding FBW7, the recognition receptor of the SCF^FBW7^ E3 ubiquitin ligase in different segments of the kidney tubular system. Deletion of *Fbxw7* in the medulla led to a juvenile-adult NPHP-like phenotype, where the decline in renal function was due to SOX9-mediated interstitial fibrosis rather than cystogenesis. In contrast, the decline of renal function in ADPKD is coupled to cystic expansion via the abnormal accumulation of FBW7 in the proximal tubules and other cell types in the renal cortex. We propose that FBW7 functions at the apex of a protein network that determines renal function in ADPKD by sensing architectural changes induced by cystic expansion.

## INTRODUCTION

Polycystic Kidney Disease encompasses a group of genetically heterogeneous diseases characterized by cystic disease and a gradual decline in renal function^1^. The most prevalent form of PKD is ADPKD with an incidence of one per 500-1,000 individuals, affecting 12.5 million people worldwide^2^. Autosomal Recessive Polycystic Kidney Disease has an incidence of one to 20,000 people^3–5^. NPHP, as a stand-alone disease, has an estimated prevalence of 1:50,000, but it has also been implicated in several syndromic forms of PKD such as Joubert, Bardet-Biedl and Meckel-Gruber syndrome^6,7^. ADTKD is an umbrella of genetic conditions with clinicopathological features that overlap with those of NPHP^8^. In addition to kidney cysts, common pathologies in syndromic forms of PKD, are progressive decline in renal function, tubulointerstitial fibrosis, and tubular atrophy or degeneration without kidney enlargement. NPHP is the most common genetic cause of renal failure in children and young adults^9,10^ and together with ADTKD account for 10-20% of children with chronic renal failure and for 1-5% of all patients undergoing dialysis or transplantation^11,12^.

While kidney function decline is the common denominator in all forms of PKD, a central question that remains poorly understood is how cystic expansion and growth induce the decline in renal function. Because cystic expansion precedes the decline in kidney function in ADPKD, it is conventionally believed that changes in kidney architecture *per se* account for the decline in kidney function. In contrast to ADPKD however, kidney function in NPHP declines at similar rates, if not faster, without excessive cystic expansion and kidney enlargement^9,10,13^. These observations suggest that the decline in renal function in different forms of PKD may be influenced by factors independent of cystic expansion.

The FBW7 recognition receptor of the Skp-Cullin-F-box (SCF)^FBW7^ E3 ubiquitin ligase processes numerous proteins for degradation via the ubiquitin-proteasome system (UPS) affecting many basic cell functions such as proliferation, metabolism, differentiation, and cell death in a cell context-dependent manner^14–16^. We have shown that FBW7 has important roles in the structure and function of the primary cilium, a central organelle in the development of PKD, where FBW7 functions as a rheostat for ciliary length influencing ciliary structure and signaling output^17,18^. Because of the wide range of proteins processed by FBW7 and its known molecular function as a component of an E3 ubiquitin ligase, we reasoned that it could be used as a discovery platform to identify protein networks underlying cystogenesis and its coupling to renal function, if deletion of *Fbxw7* in the kidney resulted in cystogenesis and a decline in renal function. The availability of reliable *in silico* tools to identify direct FBW7 targets would facilitate narrowing the search for proteins with direct roles in cystogenesis and its relationship to renal function.

Our data show that deletion of *Fbxw7* in collecting ducts resulted in a juvenile-adult NPHP-like phenotype characterized by cystogenesis, interstitial fibrosis and an early decline in renal function, which was due to fibrosis induced by the expression of SOX9, a known FBW7 target^19^. However, in the setting of ADPKD, deletion of *Fbxw7* uncoupled cystogenesis from renal function, independent of fibrosis or inflammation. This uncoupling occurred in the kidney cortex, primarily in proximal tubules, which surprisingly, did not undergo as massive cystic expansion, as seen in collecting ducts. These data lead us to propose that proximal tubules are capable of sensing non-cell autonomous architectural changes induced by cystogenesis to trigger a decline in renal function in ADPKD. This sensing mechanism involves excessive FBW7-mediated degradation of target proteins and the abnormal accumulation of several protein products of known NPHP genes functioning at the ciliary transition zone.

## RESULTS

### Deletion of *Fbxw7* in the kidney collecting ducts results in slow-progressing cysts and loss of kidney function

We deleted the *Fbxw7* gene in kidney epithelial cells using *Cdh16Cre*. This driver is active in epithelial cells of developing nephrons, the ureteric bud, and mesonephric tubules with low recombination efficiency in the proximal tubules. In the adult kidney, this promoter is active in the collecting ducts, loops of Henle, distal tubules, and to a lesser degree in proximal tubules^20^. *Cdh16Cre;Fbxw7^f/f^* mice displayed slow-progressing cystic disease without an increase in the two kidneys-to-body weight ratio (2KW/BW) and a gradual decline in kidney function (Fig. 1a-e). Cysts were present in LTA-positive proximal tubules, NKCC2-positive loop of Henle, and DBA-positive collecting ducts (Supplementary Fig. 1). Kidney function began to deteriorate as early as 1 month of age and continued to decline by 3, 7, and 10 months of age, as represented by elevated levels of serum BUN, Creatinine, and Cystatin C (Fig. 1b-d). Heterozygous *Cdh16Cre;Fbxw7^+/f^*mice showed increased levels of BUN at 7 months (Supplementary Fig. 2).

**Fig.1:**
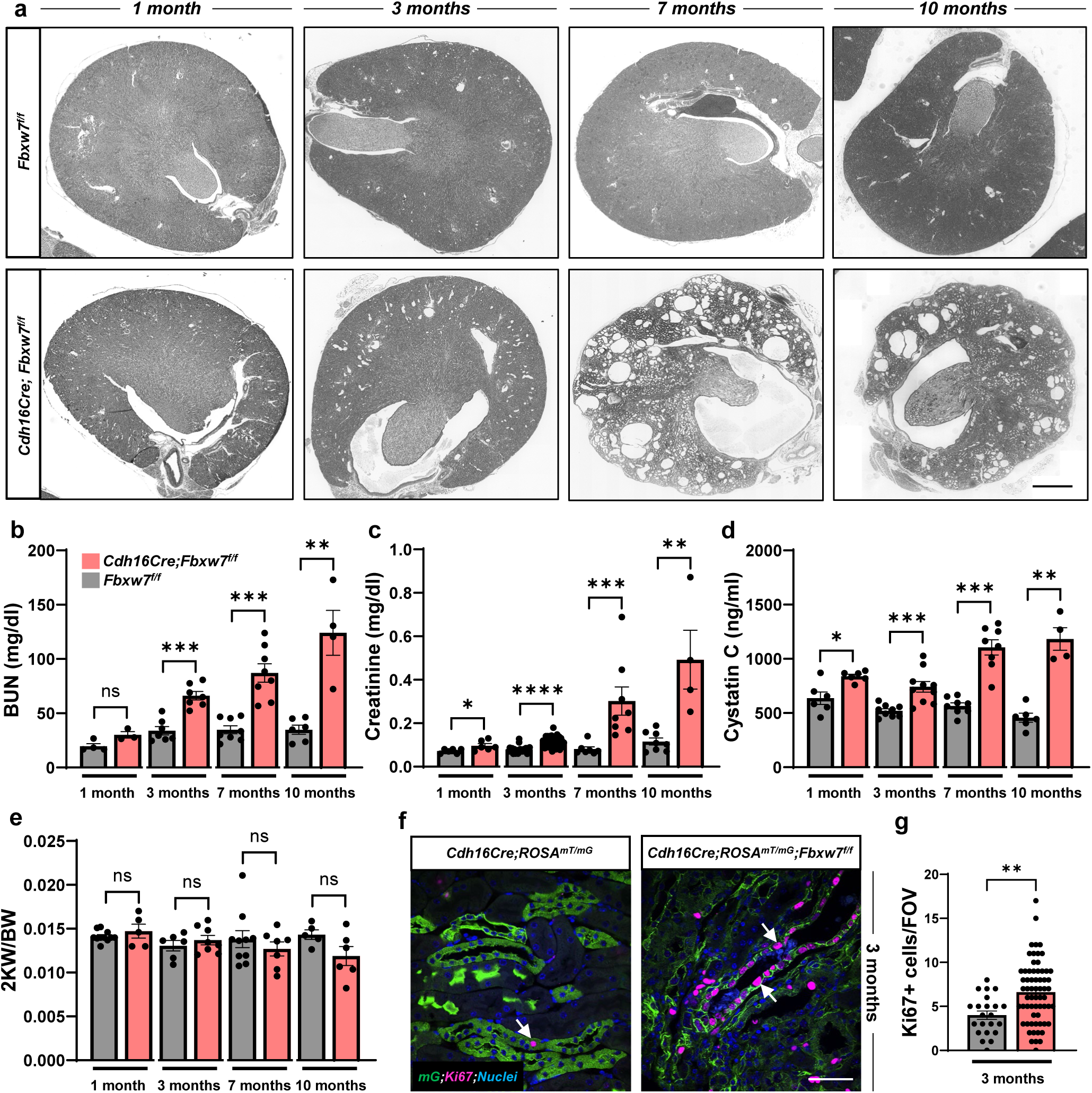
Deletion of *Fbxw7* results in a slow-progressing cyst and a decline in kidney function. (a) Representative images of whole kidney section scan showing cyst progression, (b) Serum BUN, (c) Creatinine, (d) Cytastin C, and (e) 2 kidney weight/body weight (2KW/BW) from 1-, 3-, 7-, and 10-month-old *Fbxw7^f/f^* and *Cdh16Cre;Fbxw7^f/f^*mice. Scale bar: 400 µm. Each data point represents one animal. All graphs for different time points are merged to have a common y-axis for comprehensive representation. Statistical analysis was performed using the Mann-Whitney test and is presented as the mean ± SEM (*: p<0.05, **: p<0.01, ***: p<0.001, ****: p<0.0001, ns: not significant). (f) Representative images of Ki67 staining from 3-month-old kidneys of *Cdh16Cre;ROSA^mT/mG^* and *Cdh16Cre;ROSA^mT/mG^;Fbxw7^f/f^*mice. Epithelial cells where *Cdh16Cre* is active express membrane-targeted GFP. White arrows show Ki67-positive (pink) epithelial cells where *Cdh16Cre* is active (green). Nuclei are stained with DAPI (blue). Scale bar: 50 µm. (g) Ki67 quantification. Each data point represents the Ki67-positive cells per field of view (FOV). For each genotype, n≥20 FOVs were scored from n=3 animals. Statistical analysis was performed using the Mann-Whitney test and is presented as the mean ± SEM (**: p<0.01).

We examined cyst formation and cell proliferation using *Cdh16Cre;Rosa^mT/mG^;Fbxw7^f/f^* mice that allowed us to trace the tubules affected by *Cdh16Cre*-induced recombination. From 3 months onward, cyst formation in GFP-positive tubules was accompanied by an upsurge in Ki67-positive cells, indicating increased proliferation in the *Cdh16Cre;Rosa^mT/mG^;Fbxw7^f/f^*mice (Fig. 1f-g and Supplementary Fig. 3e-f). This heightened proliferation was further supported by 5-ethynyl-2′-deoxyuridine (EdU) and phosphorylated histone H3 (phH3) staining to mark S-phase DNA synthesis and mitosis, respectively (Supplementary Fig. 3a-d). These findings are consistent with PKD in the juvenile-adult form of NPHP characterized by slow-progressing cysts and a decline in kidney function.

### Deletion of *Fbxw7* causes increased tubular cell death, thickening of the tubular basal membrane, and urine-concentrating defects

TUNEL staining of 3 and 7 months *Cdh16Cre;Rosa^mT/mG^;Fbxw7^f/f^* kidneys revealed augmented cell death (Fig. 2a-d). This rise in cell death, coupled with a reduced number of AQUAPORIN 2-positive tubules (Supplementary Fig. 4a-b), suggests a significant reduction in principal cells and degeneration of collecting ducts. Periodic acid-Schiff (PAS) staining showed disorganized architecture with tubular atrophy. Atrophic tubules had a narrow diameter, shrunken epithelium, or thickened/wrinkled basal membranes (Fig. 2e-f). *Cdh16Cre;Fbxw7^f/f^*mice began exhibiting urine concentration defects at 3 months that worsened at 7 and 10 months of age (Supplementary Fig. 4c-q). Tubular degeneration, structural changes in the tubular basement membrane, and urine-concentrating defects with acidosis are standard features of NPHP.

**Fig. 2:**
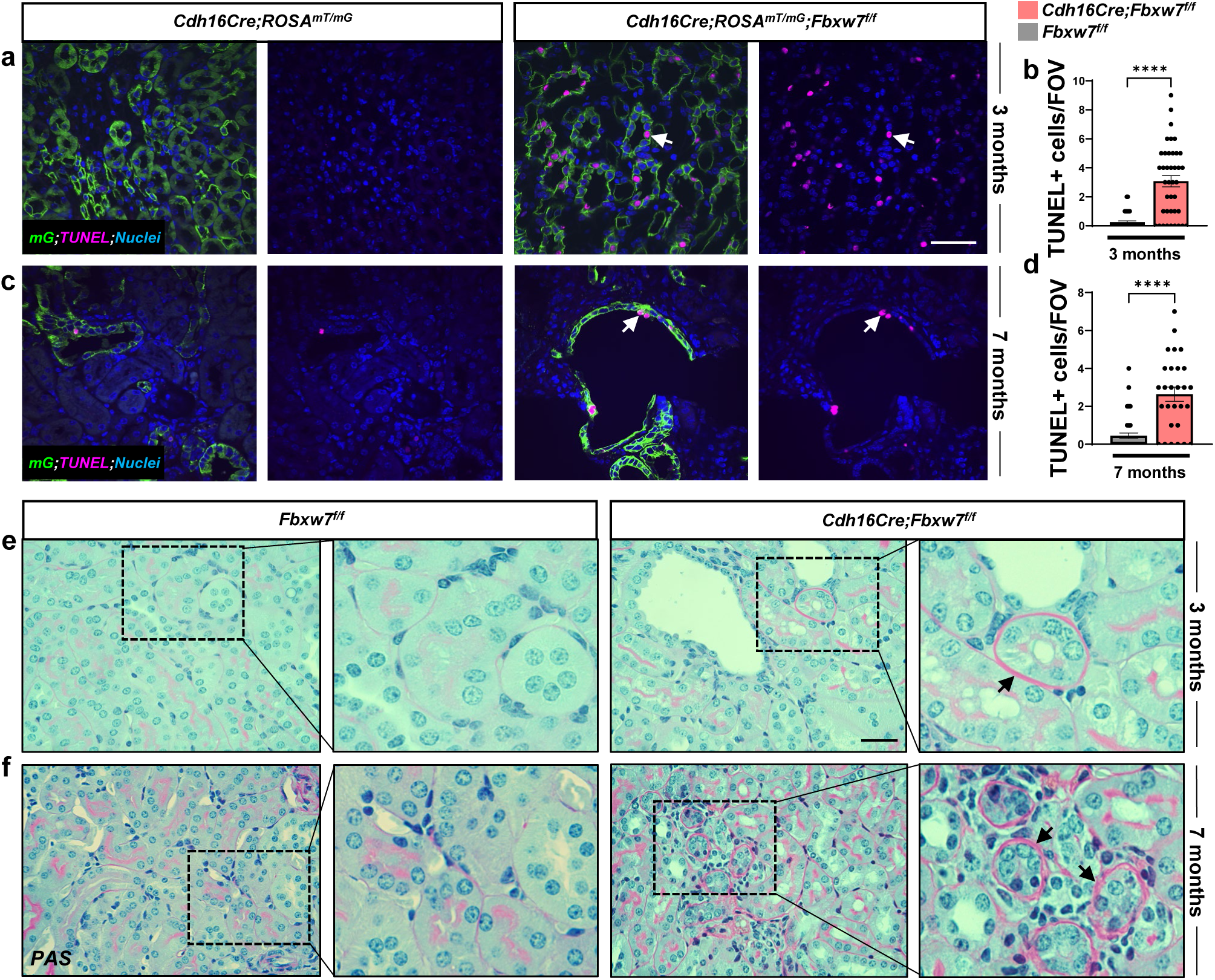
Loss of FBW7 results in increased cell death and thickening of the tubular basal membrane. (a-d) Representative images of TUNEL staining and quantification from 3- and 7-month-old kidneys of *Cdh16Cre;ROSA^mT/mG^* and *Cdh16Cre;ROSA^mT/mG^;Fbxw7^f/f^*mice. (a and c) White arrows show TUNEL-positive (pink) cells where *Cdh16Cre* is active (green). Nuclei are stained with DAPI (blue). Scale bar: 50 µm. (b and d) Each data point represents TUNEL-positive cells per FOV. For each genotype, n≥20 FOVs were scored from n=3 animals. Statistical analysis was performed using the Mann-Whitney test and is presented as the mean ± SEM (****: p<0.0001). (e-f) Representative images of PAS staining from 3- and 7-month-old kidneys of *Fbxw7^f/f^* and *Cdh16Cre;Fbxw7^f/f^* mice. Black arrows show thickened tubular basement membranes (pink). A high-magnification image of the insets is shown on the right side for each genotype. Nuclei are stained with hematoxylin (blue). Scale bar: 50 µm.

### Development of inflammation and tubulointerstitial fibrosis upon deletion of *Fbxw7*

*Cdh16Cre;Rosa^mT/mG^;Fbxw7^f/f^* kidneys displayed a marked increase in both inflammation and fibrosis, particularly evident at the corticomedullary junction from 3 months of age. Specifically, *Cdh16Cre;Rosa^mT/mG^;Fbxw7^f/f^*kidneys exhibited a significant surge in F4/80 positive macrophages near the cystic tubules, representing inflammation (Fig. 3a-b). Concurrently, there was a significant increase in Masson’s trichrome staining, depicting collagen accumulation (Fig. 3c-d). Moreover, we detected a considerable increase in VIMENTIN expression (Fig. 3e-f), typical of activated fibroblasts, and a substantial increase in α-SMOOTH MUSCLE ACTIN (αSMA), a myofibroblast marker (Fig. 3g-h), demonstrating an escalated fibrotic response in *Cdh16Cre;Rosa^mT/mG^;Fbxw7^f/f^*kidneys. These conditions worsened at 7 months of age (Supplementary Fig. 5a-d). Overall, we established that loss of FBW7 mimics NPHP pathology comprising slow-progressing cysts, tubular degeneration, inflammation, and fibrosis, all coupled with a gradual decline in kidney function.

**Fig. 3:**
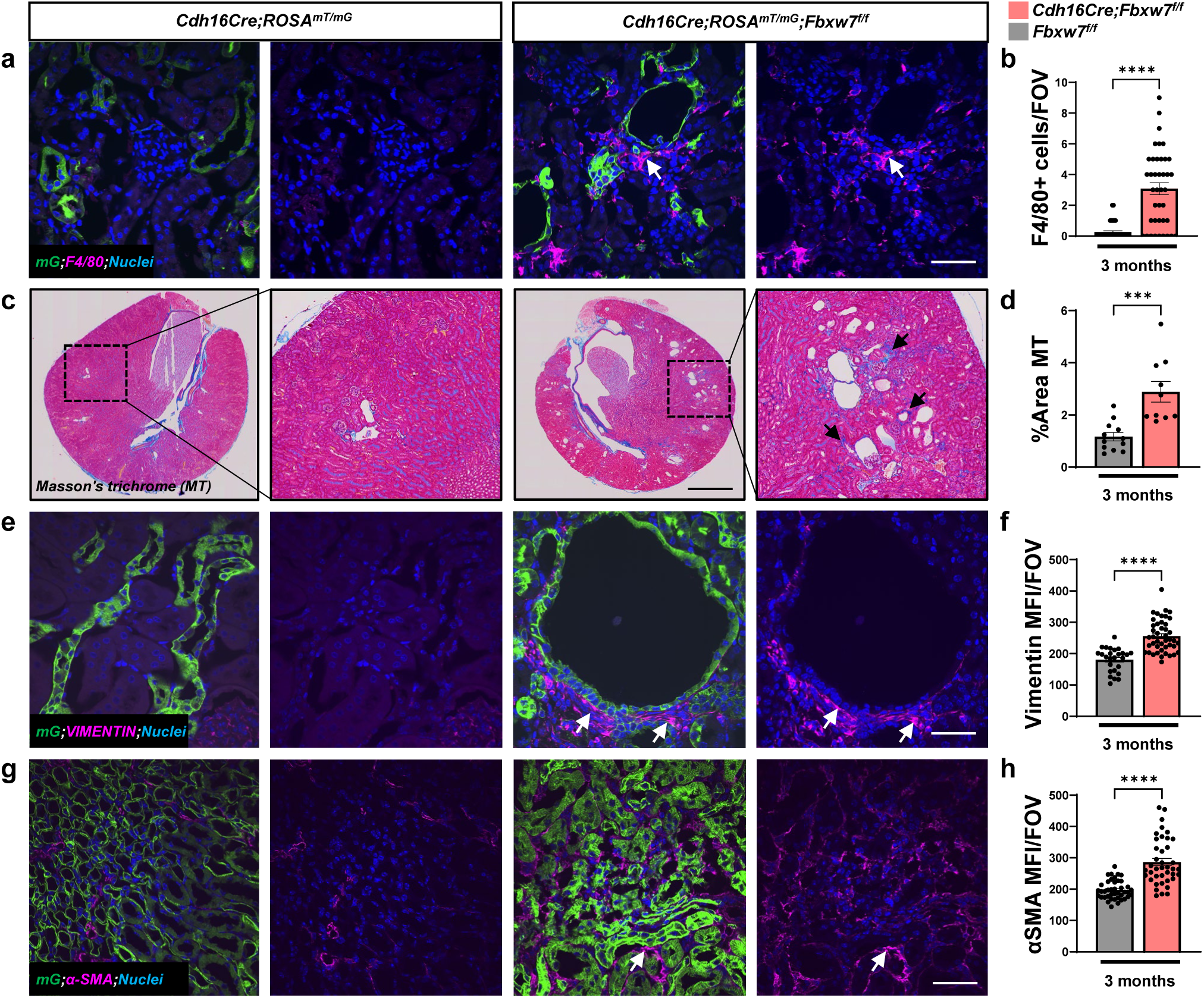
Deletion of *Fbxw7* causes inflammation and tubulointerstitial fibrosis. (a-b) Representative images and quantification of immunofluorescence staining for F4/80 from 3-month-old kidneys of *Cdh16Cre;ROSA^mT/mG^* and *Cdh16Cre;ROSA^mT/mG^;Fbxw7^f/f^* mice. (a) White arrows show F4/80-positive (pink) cells localized adjacent to the tubules where *Cdh16Cre* is active (green). Nuclei are stained with DAPI (blue). Scale bar: 50 µm. (b) Each data point represents F4/80-positive cells per FOV. For each genotype, n≥35 FOVs were scored from n=3 animals. Statistical analysis was performed using the Mann-Whitney test and is presented as the mean ± SEM (****: p<0.0001). (c-d) Representative images and quantification of Masson’s trichrome (MT) staining from 3-month-old kidneys of *Fbxw7^f/f^* and *Cdh16Cre;Fbxw7^f/f^* mice. (c) the left side image shows the whole kidney section scan and the right side image shows a high-magnification image of the insets. Black arrows show MT staining (purple) in the corticomedullary region of the kidney. Scale bar: 400 µm. (d) Each data point represents the percent area of MT staining per whole kidney section scan. For each genotype, n≥6 animals were scored. Statistical analysis was performed using the Mann-Whitney test and is presented as the mean ± SEM (***: p<0.001). (e-h) Representative images and quantification of immunofluorescence staining for (e-f) VIMENTIN and (g-h) α-SMA from 3-month-old kidneys of *Cdh16Cre;ROSA^mT/mG^* and *Cdh16Cre;ROSA^mT/mG^;Fbxw7^f/f^*mice. (e and g) White arrows show VIMENTIN or α-SMA staining (pink) adjacent to the tubules where *Cdh16Cre* is active (green). Nuclei are stained with DAPI (blue). Scale bar: 50 µm. (f and h) Each data point represents the mean fluorescence intensity (MFI) per FOV. For each genotype, (f) n≥25 or (h) n≥35 FOVs were scored from n=3 animals. Statistical analysis was performed using the Mann-Whitney test and is presented as the mean ± SEM (****: p<0.0001).

### Deletion of *Fbxw7* induced SOX9 expression in cystic tubules, and loss of *Sox9* partially ameliorates fibrosis

SOX9 is a direct target of FBW7^19^ and a pro-fibrotic factor in extra-renal tissues^21–23^. We confirmed its upregulation in renal mIMCD3 cells lacking *Fbxw7* using CRSPR/Cas9-gene editing (Fig. 4a and Supplementary Fig. 6). Since we had noted a significant increase in fibrosis around cystic tubules in the corticomedullary region in *Cdh16Cre;Rosa^mT/mG^;Fbxw7^f/f^* kidneys, we investigated SOX9 expression in these areas. We found that *Fbxw7*-null cystic tubules exhibited significant SOX9 upregulation compared to nearly undetectable levels in control animals (Fig. 4b).

**Fig. 4:**
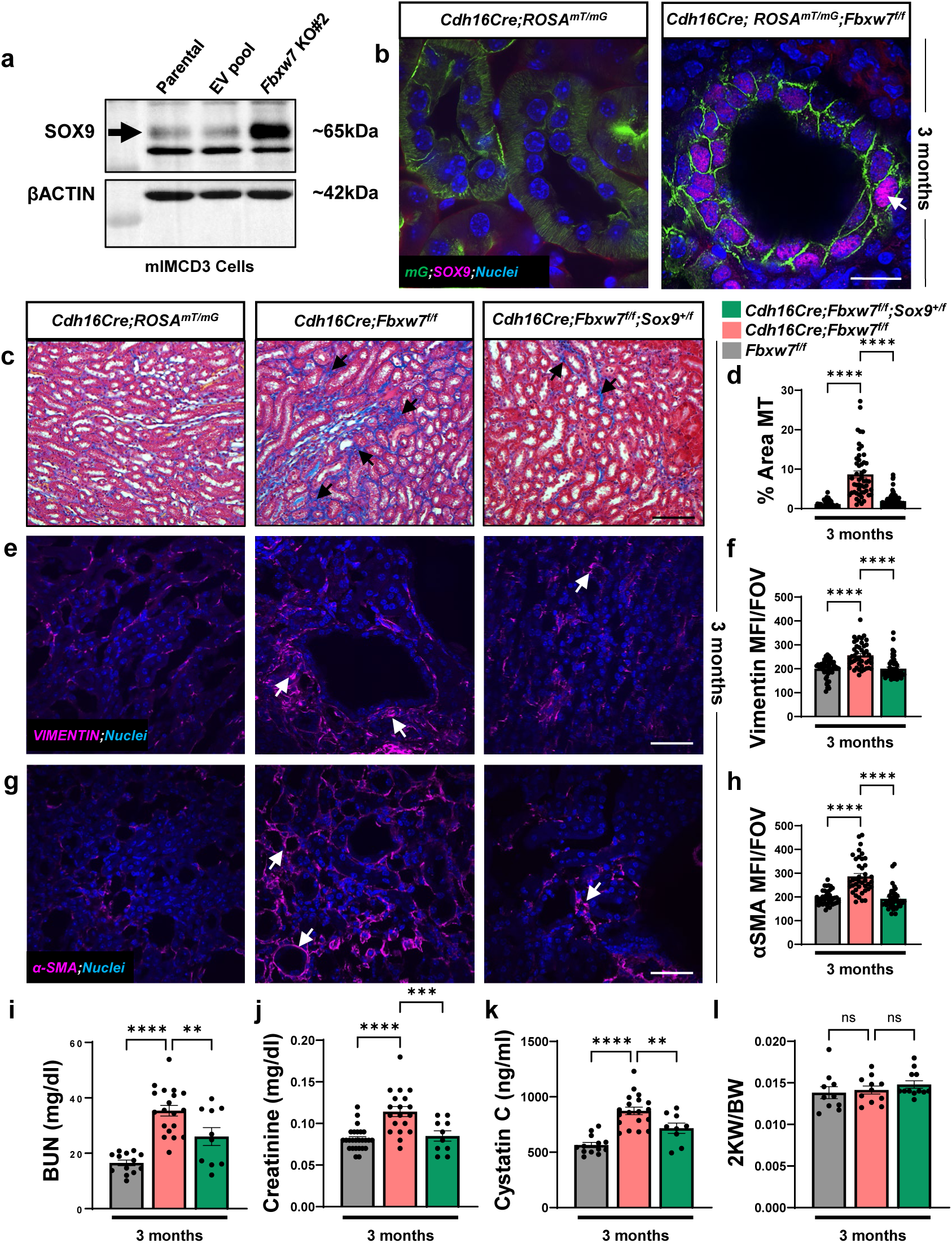
Increased expression of SOX9 contributes to tubulointerstitial fibrosis upon loss of FBW7. (a) Immunoblot showing expression levels of SOX9 from whole cell lysates of parental, empty-vector (EV) pool, and *Fbxw7*-null (*Fbxw7 KO#2*) mIMCD3 cells. (b) Immunofluorescence staining of SOX9 from 3-month-old kidneys of *Cdh16Cre;ROSA^mT/mG^* and *Cdh16Cre;ROSA^mT/mG^;Fbxw7^f/f^*mice. The white arrow shows nuclear SOX9 staining (pink) in epithelial cells where *Cdh16Cre* is active (green). Nuclei are stained with DAPI (blue). Scale bar: 20 µm. (c-h) Representative images and quantification of (c-d) MT, (e-f) VIMENTIN, and (g-h) α-SMA staining from 3-month-old kidneys of *Fbxw7^f/f^*, *Cdh16Cre;Fbxw7^f/f^*, and *Cdh16Cre;Fbxw7^f/f^;Sox9^+/f^* mice. Black arrows show (c) MT staining (purple). Scale bar: 200 µm White arrows show (e) VIMENTIN or (g) α-SMA staining (pink). Nuclei are stained with DAPI (blue). Scale bar: 50 µm. (d, f, and h) Each data point represents (d) the percent area of MT staining per FOV or (f and h) MFI per FOV. For each genotype, (d) n≥40 or (f and h) n≥35 FOVs were scored from n=3 animals. Statistical analysis was performed using one-way ANOVA followed by Šídák’s multiple comparisons test and is presented as the mean ± SEM (*: p<0.05, ***: p<0.001), ****: p<0.0001). (i) Serum BUN, (j) Creatinine, (k) Cytastin C, and (l) 2KW/BW from 3-month-old *Fbxw7^f/f^*, *Cdh16Cre;Fbxw7^f/f^*, and *Cdh16Cre;Fbxw7^f/f^;Sox9^+/f^* mice. Each data point represents one animal. Statistical analysis was performed using one-way ANOVA followed by Šídák’s multiple comparisons test and is presented as the mean ± SEM (**: p<0.01, ***: p<0.001, ****: p<0.0001, ns: not significant).

To determine whether the increased expression of SOX9 contributes to the NPHP pathology observed following *Fbxw7* deletion, we generated *Cdh16Cre;Fbxw7^f/f^;Sox9^+/f^*animals. These animals showed a decrease in fibrosis at 3 months of age, confirmed by Masson’s trichrome staining (Fig. 4c-d). Reduced fibrosis was further evidenced by decreased VIMENTIN (Fig. 4e-f) and αSMA (Fig. 4g-h) levels. Correspondingly, serum BUN, Creatinine, and Cystatin C representing kidney function were also improved with no changes in 2KW/BW in *Cdh16Cre;Fbxw7^f/f^;Sox9^+/f^*animals compared to *Cdh16Cre;Fbxw7^f/f^* animals (Fig. 4i-l). Combined wit the fact that cystogenesis is minimal in 3-month-old animals (Fig. 1a), these results suggest that fibrosis induced by the expression of SOX9 drives the decline of renal function in *Cdh16Cre;Fbxw7^f/f^* mice.

### *Fbxw7* deletion results in the downregulation of TMEM237 and causes ciliary defects

To begin probing the proteome reprogramming induced by deletion of *Fbxw7*, we employed a quantitative proteomic approach, where protein expression profiles were compared between wild type cells (parental mIMCD3 or stable pools transfected with empty vector-mock) and cells lacking FBW7 (including both mIMCD3*^Fbxw7-KO#2^* and mIMCD3*^Fbxw7-KO#7^* clones) using mass spectrometry. Of 6,975 proteins detected in 4 samples (two controls lines and two knockout lines), 57 proteins were significantly upregulated and 107 proteins were significantly downregulated. Fifteen of the 57 upregulated proteins had at least one optimal FBW7 phosphodegron and could be direct targets of FBW7. Downregulated proteins must be indirect downstream targets of FBW7. TMEM237, whose gene is mutated in patients with Joubert Syndrome (JBST14)^24^ was one of the downregulated proteins (Fig. 5a). The downregulation of TMEM237 revealed by mass spec was validated using immunoblotting of lysates of wild type and *Fbxw7*-null mIMCD3 cells (Fig. 5b-c) and kidneys (Fig. 5d). Expression of TMEM237 was drastically reduced in dilatated tubules in the corticomedullary region in 3-month-old animals (Fig. 5d). Since patients with Joubert syndrome display a renal NPHP phenotype, these data provided biochemical and mechanistic support for the NPHP-like phenotype induced by the deletion of *Fbxw7* in the renal medulla.

**Fig. 5:**
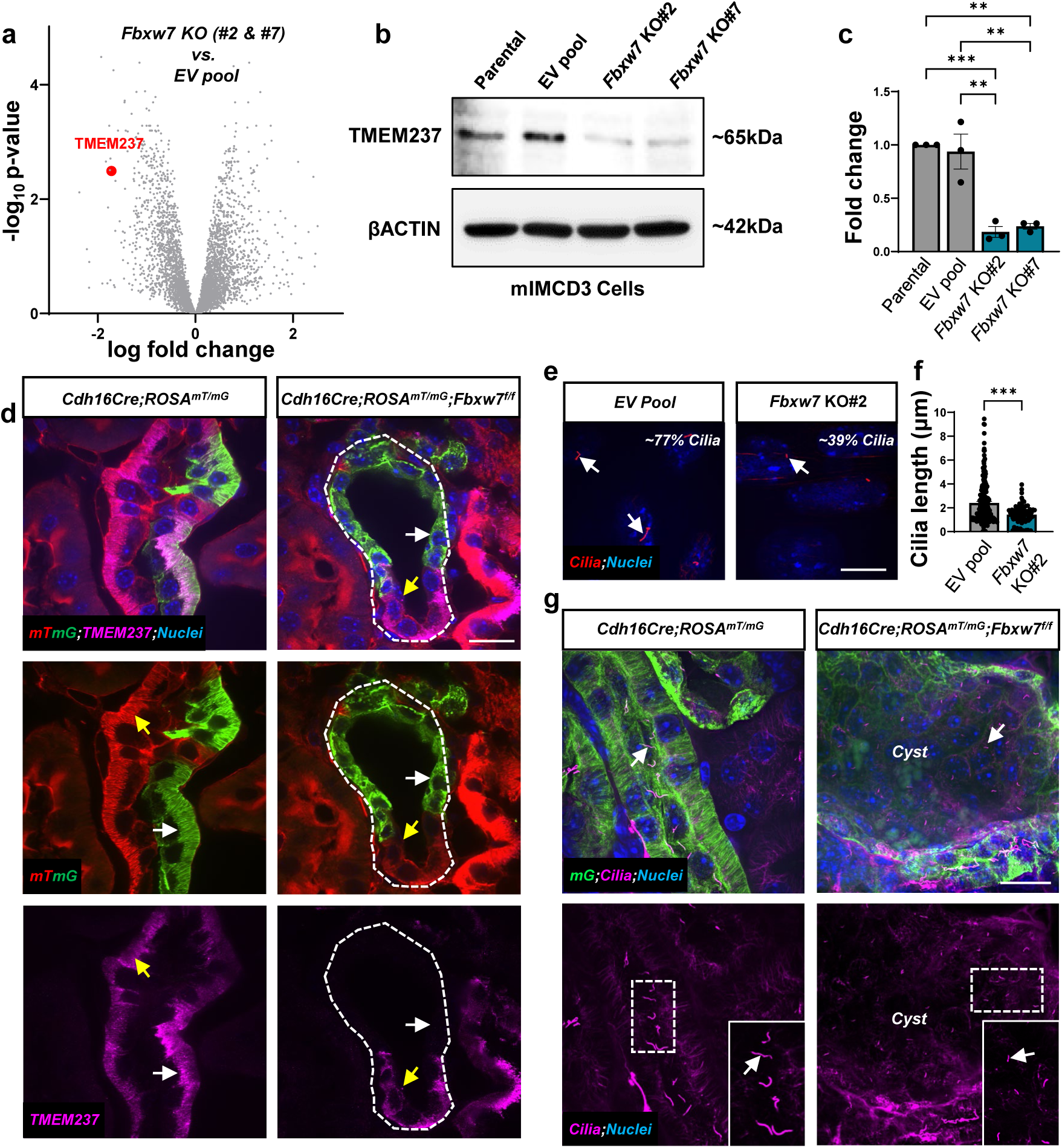
Loss of FBW7 results in TMEM237 downregulation coupled with ciliary defects. (a) Volcano plot showing differentially expressed proteins from *Fbxw7*-null cells (*Fbxw7 KO#2* and *Fbxw7 KO#7*) versus EV pool mIMCD3 cells. The red dot represents the TMEM237 protein that is significantly downregulated in *Fbxw7*-null cells compared to EV pool mIMCD3 cells. (b) Validation of quantitative proteomics using immunoblotting and (c) quantification for TMEM237 in parental, EV pool, *Fbxw7 KO#2*, and *Fbxw7 KO#7* mIMCD3 cells. Results were obtained from 3 independent experiments. (d) Representative images of immunofluorescence staining of TMEM237 from 3-month-old kidneys of *Cdh16Cre;ROSA^mT/mG^* and *Cdh16Cre;ROSA^mT/mG^;Fbxw7^f/f^*mice. The white and yellow arrows show TMEM237 expression (pink) in cells where *Chd16Cre* is active (green) or inactive (mT, red) within the same kidney tubule, respectively. Nuclei are stained with DAPI (blue). Scale bar: 20 µm. (e-f) Representative images and quantification of cilia staining from EV pool and *Fbxw7*-null (*Fbxw7 KO#2*) mIMCD3 cells. White arrows show cilia (red), and the upper right corner shows the percentage of ciliated cells. Nuclei are stained with DAPI (blue). Scale bar: 10 µm. (f) Each data point represents the ciliary length (n=3). Statistical analysis was performed using the Mann-Whitney test and is presented as the mean ± SEM (***: p<0.001). (g) Representative images of cilia staining from 3-month-old kidneys of *Cdh16Cre;ROSA^mT/mG^* and *Cdh16Cre;ROSA^mT/mG^;Fbxw7^f/f^*mice. White arrows show cilia (pink) in kidney tubules where *Cdh16Cre* is active (green). Nuclei are stained with DAPI (blue). The bottom-right corner of the lower panel shows high-magnification images of the insets. Scale bar: 10 µm.

Since the loss of most NPHP-causing genes, including *Tmem237*, is shown to cause defective cilia^24^, we further investigated if cilia were affected by the deletion of *Fbxw7*. We found that loss of FBW7 in mIMCD3 cells resulted in shorter cilia compared to that of control cells. The frequency and the length of cilia were significantly reduced in *Fbxw7*-null cells compared to that of the control cells (Fig. 5e-f). We also extended our analysis to *Cdh16Cre;Rosa^mT/mG^;Fbxw7^f/f^* kidneys, where we found that the cystic tubules displayed shorter cilia (Fig. 5g). Collectively, these findings suggest that the loss of FBW7 mimics NPHP not only phenotypically, but also mechanistically including ciliary defects and downregulation of TMEM237, a *bona fide* PKD gene.

### Loss of FBW7 rescues renal function independent of cystic expansion in an orthologous ADPKD mouse model

Genetic interactions between genes for ADPKD and syndromic forms of PKD, including NPHP and Joubert Syndrome have been hampered due to the lack of mouse models faithfully recapitulating NPHP or the renal phenotype of Joubert syndrome patients. In most NPHP types, especially in juvenile-adult types, and Joubert syndrome mouse models, renal function does not deteriorate as fast as seen in patients^25–28^. Our *Cdh16Cre;Fbxw7^f/f^* mice faithfully recapitulated juvenile-adult NPHP, allowing us to test genetic interactions with ADPKD genes. We deleted *Fbxw7*, *Pkd1*, or both using *Cdh16Cre*. *Cdh16Cre;Pkd1^f/f^*, *Cdh16Cre;Pkd1^f/f^*;*Fbxw7^+/f^*, and *Cdh16Cre;Pkd1^f/f^*;*Fbxw7^f/f^* animals displayed similar levels of cystic progression as seen by increased 2KW/BW, and no significant changes in kidney function decline as shown by serum BUN, Creatinine, and Cystatin C levels at postnatal day 16 (P16) (Supplementary Fig. 7). We further deleted *Fbxw7*, *Pkd1*, or both postnatally using tamoxifen-inducible *Cdh16Cre^ERT2^*, which also showed no changes in the 2KW/BW and decline in kidney function (Supplementary Fig. 8). Together, these results suggested that loss of FBW7 had no significant effect in ADPKD mouse models, where deletion of *Pkd1* occurs predominantly in the tubules present in the medullar region and, to a lesser extent, in the cortex region of the kidney.

Since deletion of NPHP genes result primarily in corticomedullary cysts and *Cdh16Cre;Fbxw7^f/f^* animals developed cysts in this region, along with proximal tubule cysts in the cortex (Supplementary Fig. 1a), we tested for a genetic interaction of *Fbxw7* and *Pkd1* in kidney cortex. For this purpose, we switched to the tamoxifen-inducible *UbcCre^ERT2^*-based ADPKD mouse model, where we deleted *Fbxw7*, *Pkd1*, or both postnatally. The *UbcCre^ERT2^* driver is predominantly active in the tubules present in the cortex including the corticomedullary junction with minimal, but detectable expression in the medullar region. This expression pattern is diametrically opposite to the one achieved by the *Cdh16Cre* (Supplementary Fig. 9).

*UbcCre^ERT2^;Pkd1^f/f^* animals developed massive kidney cysts in the medulla and, to a much lesser degree, in the cortex at P16, indicating that collecting ducts are extremely sensitive to the loss of *Pkd1* in forming cysts. These animals showed increased total cystic index, 2KW/BW, and a rapid decline in kidney function determined by BUN, Creatinine, and Cystatin C levels in the serum. Total cystic indices and 2KW/BW of *UbcCre^ERT2^;Pkd1^f/f^*, *UbcCre^ERT2^;Pkd1^f/f^;Fbxw7^+/f^*, and *UbcCre^ERT2^;Pkd1^f/f^;Fbxw7^f/f^*animals were similar (Fig. 6a-c). However, *UbcCre^ERT2^;Pkd1^f/f^;Fbxw7^+/f^*and *UbcCre^ERT2^;Pkd1^f/f^;Fbxw7^f/f^* animals showed almost normal kidney function based on serum BUN, Creatinine, and Cystatin C levels (Fig. 6d-f). This protective effect on kidney function upon deletion of one or two alleles of *Fbxw7* was also evident in P21 animals (Fig. 6c-f). *UbcCre^ERT2^;Fbxw7^f/f^* animals did not show any changes in the cystic index, 2KW/BW, and kidney function compared to wild-type animals at P16 (Fig. 6).

**Fig. 6:**
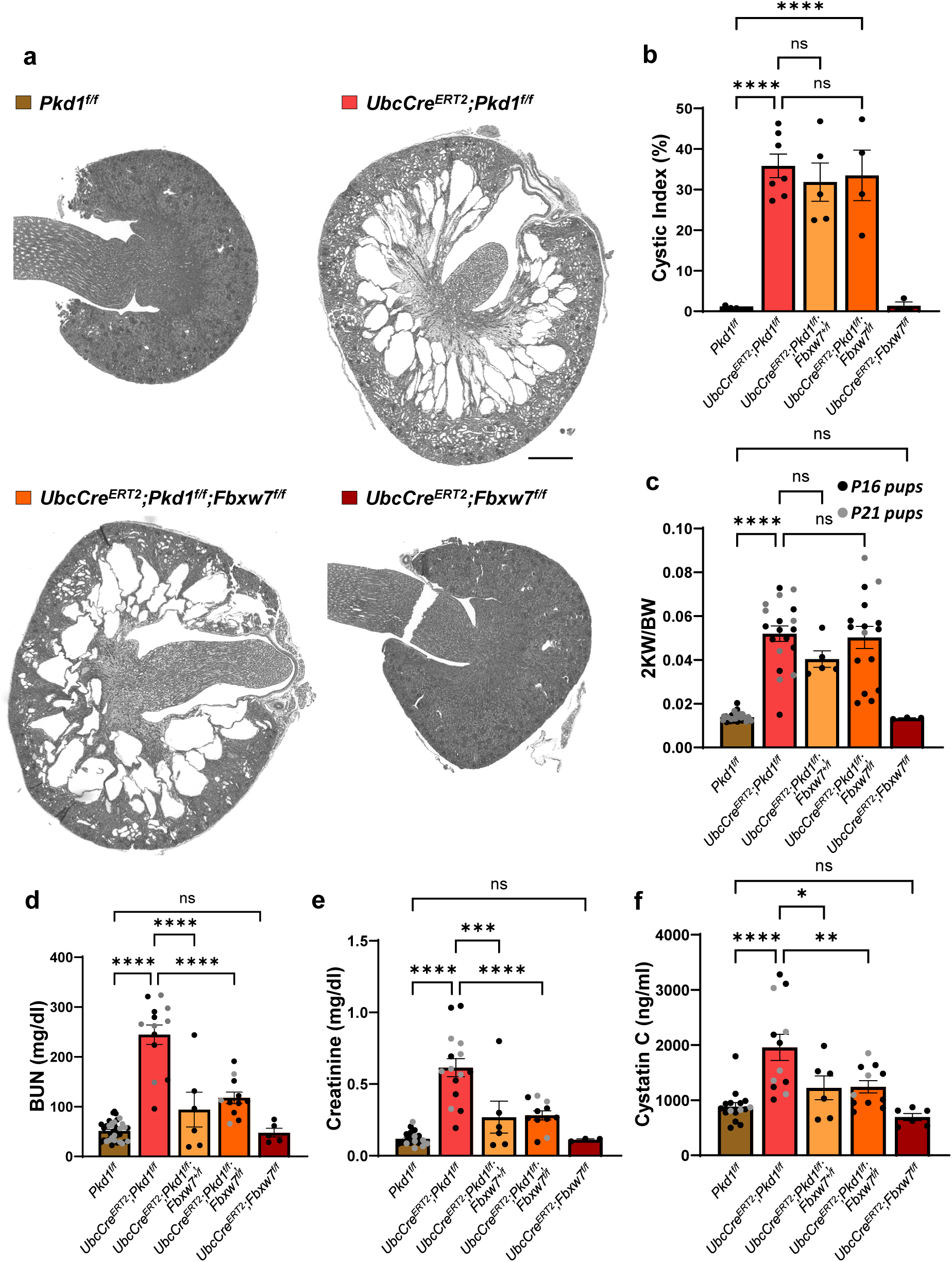
Deletion of *Fbxw7* in *UbcCre^ERT2^*-based ADPKD mouse model rescues kidney function independently of total cystic progression. (a) Representative images of whole kidney section scan showing cystic progression from *Pkd1^f/f^*, *UbcCre^ERT2^;Pkd1^f/f^*, *UbcCre^ERT2^;Pkd1^f/f^;Fbxw7^f/f^*, and *UbcCre^ERT2^;Fbxw7^f/f^*at postnatal day 16 (P16) pups. Scale bar: 400 µm. (b) Cystic index, (c) 2KW/BW, (d) Serum BUN, (e) Creatinine, and (f) Cytastin C from respective genotypes at P16 or P21. Each data point represents one animal. Black dots represent P16 pups, and grey dots represent P21 pups. Statistical analysis was performed using one-way ANOVA followed by Šídák’s multiple comparisons test and is presented as the mean ± SEM (*: p<0.05, **: p<0.01, ***: p<0.001, ****: p<0.0001, ns: not significant).

Further characterization of *UbcCre^ERT2^;Pkd1^f/f^;Fbxw7^f/f^* animals at P16 revealed no significant changes in fibrosis, cell proliferation, inflammation, or apoptosis when compared to *UbcCre^ERT2^;Pkd1^f/f^* animals (Supplementary Fig. 10). It has been shown previously that the loss of PKD1 results in longer cilia^29,30^, whereas we found that the loss of FBW7 could result in shorter cilia^17,18^. Thus, we utilized ciliary length as a phenotypic read-out assay to identify cell types of genetic interactions between *Pkd1* and *Fbxw7* across different kidney tubules. Notably, ciliary length is a sensitive measure, particularly relevant during postnatal development and in the context of ADPKD^31^. We found a significant elongation of primary cilia in epithelial cells of proximal tubules of *UbcCre^ERT2^;Pkd1^f/f^* animals, whereas *UbcCre^ERT2^;Pkd1^f/f^;Fbxw7^f/f^* animals exhibited a normalization of ciliary length. This ciliary length normalization was not evident in LTA-negative cells (Fig. 7a-b and d-e). Thus, we specifically examined the cystic index of the proximal tubules of *UbcCre^ERT2^;Pkd1^f/f^;Fbxw7^f/f^*animals. We found a partial rescue in the micro-cysts present in the proximal tubules compared to *UbcCre^ERT2^;Pkd1^f/f^* animals (Fig. 7c and f). However, this partial rescue did not significantly impact the total cystic index and 2KW/BW of *UbcCre^ERT2^;Pkd1^f/f^;Fbxw7^f/f^*animals (Fig. 6b-c). These data identified the proximal tubules as one of the cell types where PKD1 and FBW7 work antagonistically to regulate ciliary length.

**Fig. 7:**
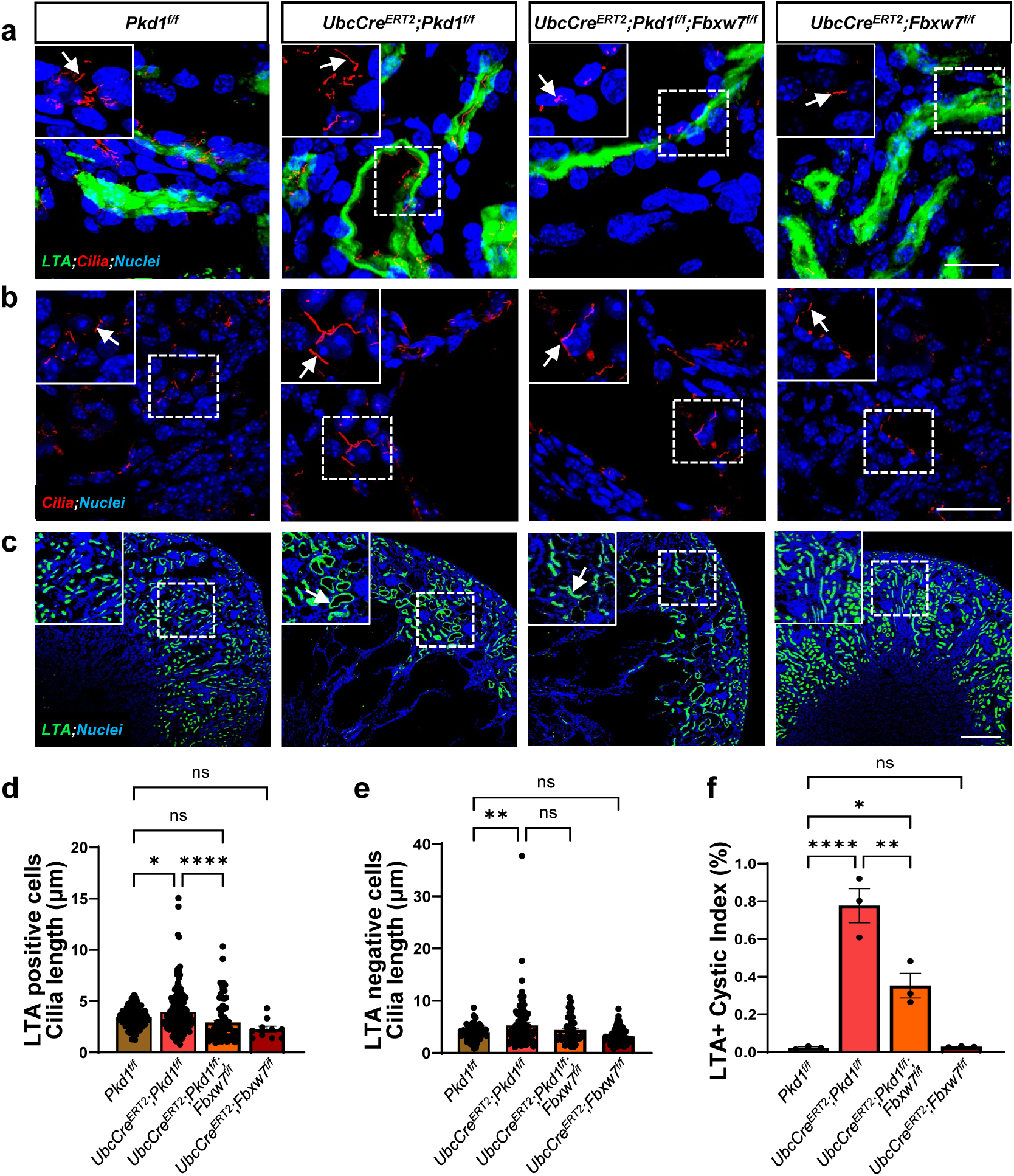
Deletion of *Fbxw7* normalizes ciliary length and ameliorates micro-cysts in proximal tubules in ADPKD. (a-c) Representative images of (a) cilia staining (red) in LTA-positive (green) or (b) LTA-negative cells, and (c) LTA-positive proximal tubules (green) with or without micro-cysts present in the kidney cortex from *Pkd1^f/f^*, *UbcCre^ERT2^;Pkd1^f/f^*, *UbcCre^ERT2^;Pkd1^f/f^;Fbxw7^f/f^*, and *UbcCre^ERT2^;Fbxw7^f/f^*pups at P16. The upper left corner shows high-magnification images of the insets, and the white arrows show (a-b) cilia staining (red) or (c) proximal tubules (green). Nuclei are stained with DAPI (blue). Scale bar: (a-b) 20 µm and (c) Scale bar: 200 µm. (d-f) Quantification of ciliary length in (d) LTA-positive cells, (e) LTA-negative cells, and (f) LTA-positive cystic index from respective genotypes. Each data point represents (d-e) the length of cilia per cell or (f) the LTA-positive cystic index per animal (n=3). For each genotype, (d) n≥10 or (e) n≥25 FOVs were scored from n=3 animals. Statistical analysis was performed using one-way ANOVA followed by Šídák’s multiple comparisons test and is presented as the mean ± SEM (*: p<0.05, **: p<0.01, ****: p<0.0001, ns: not significant).

### Upregulation of FBW7 and other NPHP genes in ADPKD tissues

Expression analysis revealed a significant upregulation of *Fbxw7* mRNA (Fig. 8a-b) and protein levels (Fig. 8c-e) in the kidney, especially in the proximal tubules of *UbcCre^ERT2^;Pkd1^f/f^*animals (Fig. 8a-b). This FBW7 upregulation was also apparent in kidney sections obtained from 3 out of 3 ADPKD patients (Fig. 8f). To determine whether there was a pattern including other NPHP genes, we examined expression levels of NPHP1 and NPHP4 in ADPKD patient biopsies. We observed a significant upregulation of NPPH1 and NPHP4 in 3 out of 3 ADPKD patients (Supplementary Fig. 11). These data show the upregulation of FBW7 and other NPHP genes in mouse *Pkd1*-null and human ADPKD kidneys.

**Fig. 8:**
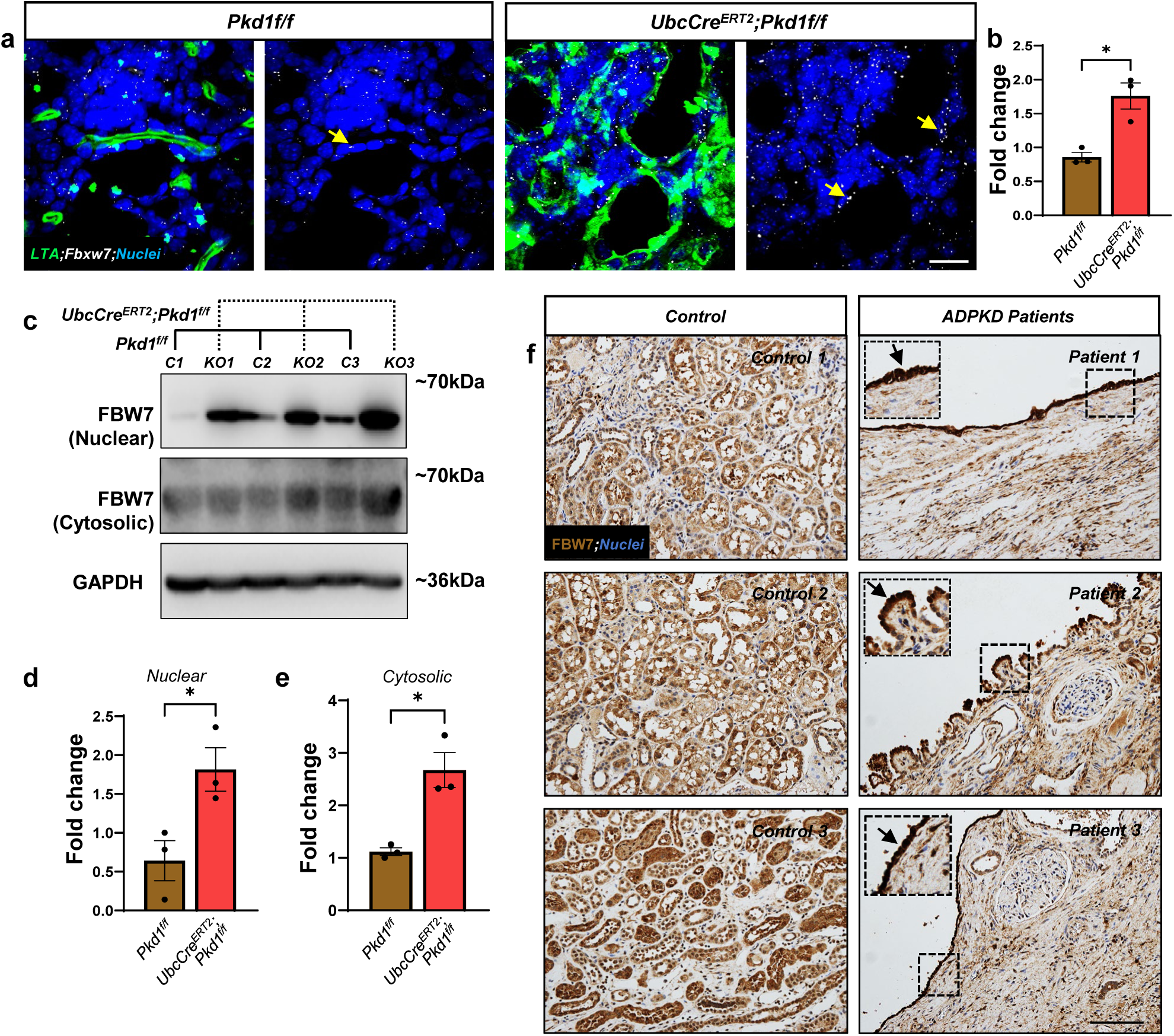
Deletion of *Pkd1* increases *Fbxw7* mRNA and FBW7 protein. (a-b) Representative images and quantification of RNAScope for *Fbxw7* transcript levels in LTA-positive proximal tubules from *Pkd1^f/f^* and *UbcCre^ERT2^;Pkd1^f/f^*pups at P16. Yellow arrows show *Fbxw7* expression (white) in LTA-positive proximal tubules (green). Nuclei are stained with DAPI (blue). Scale bar: 20 µm. (b) Each data point represents a fold change in the MFI per FOVs. For each genotype, n≥3 FOVs were scored from n=3 animals. Statistical analysis was performed using the Mann-Whitney test and is presented as the mean ± SEM (*: p<0.05). (c-e) Immunoblots showing (c) expression levels and (d-e) quantification of FBW7 in nuclear and cytoplasmic fractions from whole kidney lysates of *Pkd1^f/f^* (n=3, labeled as C1, C2, and C3) and *UbcCre^ERT2^;Pkd1^f/f^*(n=3, labeled as KO1, KO2, and KO3) P16 pups. (f) Representative images of FBW7 expression in normal (n=3) and ADPKD patients (n=3) kidneys using immunohistochemistry. The upper left corner shows a high-magnification image of the insets, and the black arrows show FBW7 expression in the cystic epithelium. Scale bar: 1000 μm.

## DISCUSSION

A fundamental, yet mechanistically, poorly understood question is whether and how cystic expansion triggers the decline in renal function in PKD. Our study identifies FBW7 as a crucial mediator modulating renal function in ADPKD and NPHP in a cell/tubule-specific manner. We show that deletion of *Fbxw7* in the kidney medulla captures the classic triad of renal pathology of NPHP, which are corticomedullary cysts-interstitial fibrosis-tubular degeneration, accompanied by a gradual decline in renal function. In this setting, SOX9-induced fibrosis has a dominant role in driving the decline in renal function. In the setting of ADPKD however, FBW7 is abnormally accumulated in the proximal tubules in the cortex and its heterozygous deletion uncouples cystogenesis from a decline in renal function. Deletion of *Fbxw7* in the medulla does not uncouple cystic expansion from the decline in renal function. These data lead us to propose a model whereby deletion of *Pkd1* in the kidney cortex induces the expression of FBW7 that signals the decline in renal function. Our data have important implications in understanding the mechanisms that control renal function in dominant and syndromic forms of PKD. They go a step further to help us understand how cystogenesis is reciprocally linked to renal function in ADPKD. This is especially relevant in developing and/or evaluating therapeutic approaches to ADPKD, where suppressing cystic growth and effects on renal function should be considered equally important.

We undertook an approach of proteome reprogramming via the deletion of *Fbxw7* in the collecting duct system to identify protein networks responsible for cystogenesis. We chose the collecting ducts because they are very sensitive to cystogenesis. Our approach led to the development of a mouse model of slowly progressing corticomedullary cystogenesis along with fibrosis and tubular degeneration, the classic triad of renal pathology in NPHP. Using a quantitative proteomics approach, we identified TMEM237, whose gene is inactivated in patients with Joubert syndrome^24^, as an indirect target of the deletion of *Fbxw7*. This is consistent with the fact that renal pathology of patients with Joubert syndrome is closely related to NPHP^24^, suggesting the reduction of TMEM237 levels in the collecting system may contribute to the NPHP-like phenotype in these mice. Aided by the ability to identify direct FBW7 targets, we show that the upregulation of SOX9 is the driver of the decline in renal function in this NPHP-like mouse model by promoting interstitial fibrosis. The functional role of SOX9 in renal function in NPHP-like pathology or any fibrocystic disease within the PKD spectrum is shown here for the first time. Collectively, these data identify the FBW7-mediated degradation of SOX9 as a functional pathway in controlling renal function in an NPHP-like pathology via interstitial fibrosis and SOX9 as a potential pharmacological target for the treatment of other fibrocystic diseases, which is currently lacking.

Deletion of *Fbxw7* in the proximal tubules in the renal cortex did not produce an obvious kidney phenotype, but it did have a strong modifier role in controlling renal function in ADPKD. These data highlight the importance of proximal tubules in preserving renal function in ADPKD and raise the possibility that PKD1 may work antagonistically to FBW7, TMEM237, NPHP1, NPHP4, and possibly other NPHP gene products in the proximal tubules to regulate renal function independently of their roles in cystogenesis. Because NPHP gene products are known to have important functions in forming and maintaining the structural integrity of the ciliary transition zone^32^, a domain that controls the trafficking of integral and soluble proteins in and out of primary cilia, it is tempting to speculate that the transition zone may mediate the protective effect in renal function. While loss-of-function studies have solely supported the involvement of NPHP gene products in the transition zone, our study suggests that excessive accumulation of these proteins may cause structural and/or functional changes in the transition zone in proximal tubule cells, enabling these cells to sense the architectural changes in collecting ducts and other parts of the kidney. Considering that cilia ablation can suppress cystic growth and rescue kidney function in ADPKD mouse models^33^, it is plausible that the coupling between cystic growth and renal function may be mediated by a subset of specific functions from the transition zone.

Deletion of *Fbxw7* in *Pkd1/Fbxw7* mice shows that massive kidney cysts in the medulla can have little effect on overall kidney function. However, how can these results reconcile with the lack of effect of the deletion of *Fbxw7* in *Cdh16Cre;Pkd1^f/f^;Fbxw7^f/f^*mice and wealth of data showing a decline in renal function in animal models where deletion of genes responsible for the prevention of cystogenesis is restricted to collecting ducts? One possibility is that other parts of the tubular system have a similar sensing mechanism, as the proximal tubules, but with a much higher threshold (Supplemental Fig. 12). In this scenario, renal function will eventually decline in *UbcCre^ERT2^;Pkd1^f/f^;Fbxw7^f/f^*kidneys, but at a later time point. Because proximal tubular cysts were much smaller in size and number compared to the ones from the collecting ducts, yet coupling seemed to occur primarily in proximal tubules, we favor the idea that there is a reciprocal relationship between thresholds for cystogenesis and thresholds for coupling of cystogenesis and a decline in renal function. We propose that cystogenesis threshold decreases, whereas coupling threshold increases from the cortex to the medulla (Supplemental Fig. 12). Both thresholds are set by normally functioning PKD1. Interestingly, the only known gradient with increasing intensity from the cortex to the medulla is the osmolarity gradient^34^, raising the possibility that osmolarity may drive cyst growth and expansion in *Pkd1*-null kidneys, but not the coupling between cystogenesis and renal function. PKD1 could be part of the apparatus that senses osmolality and/or mechanotransduction. Whether these two sensory modalities are separate, or one is unclear.

Our data show that the coupling between cystic expansion and the decline in renal function occurs primarily in proximal tubules. As discussed above, it appears that the collecting ducts are much more sensitive to the deletion of *Pkd1* in forming cysts compared to proximal tubules, yet renal function is determined by the proximal tubules. This observation prompts us to consider the following possibility. PKD1 in proximal tubules, in addition to its role to control kidney tubule diameter, has a separate function to sense cell-autonomous (i.e., proximal tubules) and non-cell-autonomous (i.e., collecting ducts) architectural changes and trigger a decline in renal function. This second function culminates in the upregulation of FBW7 and other NPHP genes. While the underlying mechanisms of the upregulation of FBW7 in *Pkd1*-mutant kidneys are unknown, several possibilities exist. First, deletion of *Pkd1* causes an upregulation of p53, which has been shown to transcriptionally activate *Fbxw7* gene expression^30,35^. Second, deletion of *Pkd1* leads to the upregulation of USP28, a known deubiquitinase of FBW7^30,36^. Third, upregulation of the kinase(s), such as GSK3α/β, CDK1, CDK5, and possibly others that prime FBW7 relevant targets could account for its ability to degrade targets connecting cystic growth to renal function^37–39^. Interestingly, CDK1/2/5 levels and/or activities have been shown to be induced in various forms of PKD^37,38,40^. Overall, it is reasonable to suggest that the root cause of coupling cystic expansion and a decline in renal function could be due to the induced expression/activity of FBW7 secondary to cellular stress induced by metabolic reprogramming, centrosomal defects, and/or mechanical distension of the tubular cells^16^. It is also possible that the mechanisms of the upregulation of other NPHP genes are independent from the mechanism that involves the upregulation of FBW7 and the upregulation of other NPHP genes have a synergistic role in the coupling mechanism. However, because we see an almost complete rescue of renal function in *UbcCre^ERT2^;Pkd1^f/f^;Fbxw7^f/f^* mice, we favor the idea that FBW7 functions upstream of the other upregulated NPHP genes.

Our rationale for examining expression levels of other NPHP genes in *Pkd1*-null cells stemmed from the data showing that deletion of *Fbxw7* resulted in an NPHP-like phenotype. We observed a pattern, whereby NPHP1 and NPHP4 which form a protein nodule in the ciliary transition zone were also upregulated. Normalization of FBW7 levels restored renal function without a measurable effect on overall cystic index. Because disease progression in people with ADPKD and NPHP1 is milder compared to NPHP1 alone^41^, we speculate that similar effects will be seen in compound *Pkd1/Nphp1*, *Pkd1/Nphp4*, or *Pkd1/Tmem237* mice.

Clinically, *FBXW7* mutations have been documented in a wide range of human cancers^16,42^. Thus, exploring the presence of *FBXW7* mutations in patients with ADPKD or NPHP is an intriguing prospect. Given the high prevalence of naturally occurring *PKD1/PKD2* mutations in ADPKD and *FBXW7* mutations in several types of cancers^14,16,42^, it is reasonable to speculate that compound heterozygous mutations in *PKD1/PKD2* and *FBXW7* genes may be present in a subset of ADPKD patients. According to our study, renal function should decline slower in these individuals and thus, they may belong to a group of ADPKD patients that do not show severe disease symptoms. We predict the opposite clinical picture for NPHP patients. NPHP patients with either heterozygous or homozygous mutations in genes like *NPHP1*, *NPHP2*, or NPHP4, and an additional mutated *FBXW7* allele should exhibit exacerbated NPHP pathology, potentially categorizing FBW7 as a disease modifier in NPHP. This possibility is supported by heterozygous *Fbxw7* mice which show a trend for a decline in renal function in 7-month-old mice. Because global deletion of *Fbxw7* leads to embryonic lethality in mice, we do not expect to identify humans with equivalent homozygous *FBXW7* mutations.

Overall, we show for the first time that loss of FBW7 not only mimics NPHP pathology but also determines kidney function in ADPKD without significantly impacting total cyst growth and kidney enlargement. These effects are explained by deletion versus overexpression of FBW7, respectively, in specific kidney segments. Our findings warrant a thorough re-evaluation of the traditional view that cystogenesis drives kidney function decline and accentuates the divergent roles of tubules in the medulla versus the cortex in separating cystic growth from kidney function.

## METHODS

### Mouse models, tissue processing, and serum analysis

All procedures were performed according to the requirements of the Institutional Animal Care and Use Committee. Mouse strains used were *Cdh16Cre, Cdh16Cre^ERT2^, UbcCre^ERT2^, Fbxw7^f/f^*, *Pkd1^f/f^*, *Sox9^f/f^*, *and Rosa^mT/mG^*. All mice were on C57BL/6-background. Both sexes were used in all studies. For experiments with *Cdh16Cre*, animals were collected at P16, 1-, 3-, 7-, and 10-month time points. Metabolic cages were used for urine collection over a period of 16 hours. For experiments with *Cdh16Cre^ERT2^* and *UbcCre^ERT2^*, nursing dams were intraperitoneally injected with 4-hydroxytamoxifen (4-OHT; Sigma-Aldrich Cat. no. H6278) diluted in corn oil, at 100 μg/gram of nursing dam body weight, from postnatal days 2–6 (P2–P6) as described^30^. Pups were collected either at P16 or P21. For all experiments, kidneys and body weights were measured. The kidneys were later cut transversally and then fixed in either 10% neutral buffered formalin for paraffin embedding and sectioning or 4% paraformaldehyde for OCT embedding and cryo-sectioning. Tissue processing, embedding, sectioning, and histology stainings were done at the Cell Biology Histology Core and the Stephenson Cancer Center Histology Core of the University of Oklahoma Health Sciences Center (OUHSC). Cystic index and Masson’s trichrome staining quantification were done using ImageJ. Blood was collected via cardiac puncture, and serum was collected using serum separator tubes (BD Microtainer 365967). Serum BUN and Cystatin C analysis were done using Invitrogen BUN Colorimetric Detection Kit (Cat. no. EIABUN) and R&D Systems Cystatin C Quantikine ELISA Kit (Cat. no. MSCTC0), respectively. Serum creatinine analysis was done using Liquid chromatography–mass spectrometry (LC-MS) by the Bioanalytical Core, O’Brien Center for Acute Kidney Injury Research at the University of Alabama at Birmingham.

### Generation of Fbxw7-null mIMCD3 cells

mIMCD3 cells underwent transfection using a lentivirus carrying a vector that encodes Cas9. This vector was either an empty vector or included a single guide RNA (sgRNA) specifically targeting *Fbxw7* (sequence: 5′-CACCGATGAAGTCTCGCTGGAACTG-3′). The chosen sgRNA sequence and corresponding vector were previously validated for their efficiency in knocking out *Fbxw7* in mouse cell lines^18^. Following 24 hours post-transduction, the cells were subjected to puromycin treatment at a concentration of 2 µg/ml for 48 hours. Subsequently, to initiate single-cell-based colonies, the cells were dissociated with trypsin and individually sorted into 96-well plates using FACS. Several single-cell-based clones were then expanded and assessed for the absence of FBW7 through Western blot analysis. We successfully identified two clones with complete elimination of FBW7 protein, *Fbxw7*-KO #2 and *Fbxw7*-KO#7. Both parental mIMCD3 cells and those stably transfected with the empty vector were used as controls for comparative purposes.

### Immunofluorescence, TUNEL, RNAScope, and Microscopy

For immunofluorescence studies, 10 μm or 50 μm kidney cryosections or mIMCD3 cells serum starved for 48 hours on glass coverslips were used. Sections were washed 3 times with PBS, 5 minutes each, and then incubated in blocking solution (3% B.S.A., 0.15% Triton X-100, 5% Goat Serum in PBS) for 1 hour at room temperature. Samples were then incubated in primary antibody overnight at 4°C, washed with PBS 3 times 5 minutes each, and then incubated with the appropriate secondary antibodies in a blocking solution for 1 hour at room temperature. Sections were then washed with PBS 3 times for 5 minutes each, mounted with Diamond DAPI, and visualized with confocal microscopy. Antibodies used were acetylated-α-tubulin (611b, 1:2,000; Sigma-Aldrich), Ki67 (1:200; ab15580), PE-conjugated F4/80 (1:50; BD Pharmingen™), NKCC2 (1:200, ThermoFisher PA5142445), Phospho-HISTONE H3 (1:200, Cell Signaling 9701), VIMENTIN (1:200, Cell Signaling 5741), α-SMOOTH MUSCLE ACTIN (1:200, Cell Signaling 19245), AQUAPORIN 2 (1:200, ThermoFisher PA5-78808 ), SOX9 (1:200, Sigma-Aldrich AB5535), and TMEM237 (1:200, ThermoFisher PA5-63013). Fluorescein (1:1000; Vector Laboratories FL-1321) or Cy5 (1:1000; USBIO 518482) conjugated Lotus Tetragonolobus Lectin (LTA) and Rhodamine conjugated Dolichos Biflorus Agglutinin (DBA) (1:1000; Vector Laboratories RL-1032-2), was added to co-stain proximal tubules and collecting ducts, respectively, in the secondary antibody solution wherever needed. For EdU incorporation, mice received 50 mg per kg of body weight of EdU (Invitrogen, catalog no. A10044) by intraperitoneal injection 4 hr before euthanasia. EdU staining was done using Click-iT EdU Alexa Fluor-594 Imaging Kit (Invitrogen, catalog no. C10339). For TUNEL staining, the One-step TUNEL Apoptosis Kit was used per the manufacturer’s instructions (Elabscience E-CK-A324). *Fbxw7* mRNA expression was determined using a pre-designed probe (Cat No. 420371-C3) and RNAscope® Multiplex Fluorescent kit from Advanced Cell Diagnostics per the manufacturer’s instructions. All images were obtained either by an Olympus FV1000 confocal microscope, Nikon CSU-W1 SoRa Spinning Disk confocal microscope, or Olympus VS120 Virtual Slide System.

### Human ADPKD Samples

Normal and ADPKD tissue section slides were obtained from the Kansas PKD Research and Translational Core Center (U54DK126126) and the National PKD Research Resource Consortium. Samples were fixed in 4% P.F.A. overnight, then rinsed and stored in 70% EtOH until blocked in paraffin. Immunohistochemistry for FBW7 (1:1600; Bethyl antibody A301-721A), NPHP1 (1:200, ThermoFisher PA5-78808 ), and NPHP4 (1:50, ThermoFisher PA5-78808 ) was performed on 5 µm-thick paraffin sections by the Stephenson Cancer Center Histology Core at The University of Oklahoma Health Sciences Center (OUHSC).

### Immunoblotting

Frozen kidneys from P16 pups were lysed using a plasma membrane protein extraction kit (101Bio P503L). Cytoplasmic and nuclear fraction was used for the Western blots. The antibodies used were FBW7 (1:1000 in a 1:1 ratio, Bethyl A301-720A, and A301-721A) and GAPDH (1:4,000; Genetex). mIMCD3 cell lysates were made using RIPA lysis buffer. For FBW7 blots from mIMCD3 cells, abnova FBW7 antibody (1:100, abnova H00055294-B01P) was used to pull down FBW7 and Bethy FBW7 antibodies (1:1000 in a 1:1 ratio, Bethyl A301-720A, and A301-721A) were used to detect FBW7 levels in controls and *Fbxw7*-null single cell clones. Other antibodies used were SOX9 (1:1000, Sigma-Aldrich AB5535), TMEM237 (1:1000, Atlas antibodies HPA052596), and β-ACTIN (1:1000, sc-47778). Densitometric quantification was performed with ImageJ.

### Proteomic screen in mIMCD3 cells

Total protein from each sample was reduced, alkylated, and purified by chloroform/methanol extraction prior to digestion with sequencing grade modified porcine trypsin (Promega). Tryptic peptides were then separated by reverse phase XSelect CSH C18 2.5 um resin (Waters) on an in-line 150 x 0.075 mm column using an UltiMate 3000 RSLCnano system (Thermo). Peptides were eluted using a 60 min gradient from 98:2 to 65:35 buffer A:B ratio. Eluted peptides were ionized by electrospray (2.2 kV) followed by mass spectrometric analysis on an Orbitrap Exploris 480 mass spectrometer (Thermo). To assemble a chromatogram library, six gas-phase fractions were acquired on the Orbitrap Exploris with 4 m/z DIA spectra (4 m/z precursor isolation windows at 30,000 resolution, normalized AGC target 100%, maximum inject time 66 ms) using a staggered window pattern from narrow mass ranges using optimized window placements. Precursor spectra were acquired after each DIA duty cycle, spanning the m/z range of the gas-phase fraction (i.e. 496-602 m/z, 60,000 resolution, normalized AGC target 100%, maximum injection time 50 ms). For wide-window acquisitions, the Orbitrap Exploris was configured to acquire a precursor scan (385-1015 m/z, 60,000 resolution, normalized AGC target 100%, maximum injection time 50 ms) followed by 50x 12 m/z DIA spectra (12 m/z precursor isolation windows at 15,000 resolution, normalized AGC target 100%, maximum injection time 33 ms) using a staggered window pattern with optimized window placements. Precursor spectra were acquired after each DIA duty cycle. To assemble a data-dependent spectral library, a pooled peptide reference samples was separated into 46 fractions on a 100 x 1.0 mm Acquity BEH C18 column (Waters) using an UltiMate 3000 UHPLC system (Thermo) with a 50 min gradient from 99:1 to 60:40 buffer A:B ratio under basic pH conditions, then consolidated into 18 super-fractions. Each super-fraction was further separated by reverse phase XSelect CSH C18 2.5 um resin (Waters) on an in-line 150 x 0.075 mm column using an UltiMate 3000 RSLCnano system (Thermo). Peptides were eluted using a 60 min gradient from 98:2 to 65:35 buffer A:B ratio. Eluted peptides were ionized by electrospray (2.2 kV) followed by mass spectrometric analysis on an Orbitrap Exploris 480 mass spectrometer (Thermo). MS data were acquired using the FTMS analyzer in profile mode at a resolution of 120,000 over a range of 375 to 1500 m/z. Following HCD activation, MS/MS data were acquired using the FTMS analyzer in centroid mode at a resolution of 15,000 and normal mass range with normalized collision energy of 30%. Buffer A = 0.1% formic acid, 0.5% acetonitrile. Buffer B = 0.1% formic acid, 99.9% acetonitrile. Both buffers adjusted to pH 10 with ammonium hydroxide for offline separation. Following data acquisition, data were searched using an empirically corrected library against the UniProt Mus musculus database (April 2022) and a quantitative analysis was performed to obtain a comprehensive proteomic profile. Proteins were identified and quantified using EncyclopeDIA^43^ and visualized with Scaffold DIA using 1% false discovery thresholds at both the protein and peptide level. Protein exclusive intensity values were assessed for quality using ProteiNorm^44^. The data was normalized using cyclic loess and statistical analysis was performed using linear models for microarray data (limma) with empirical Bayes (eBayes) smoothing to the standard errors^45^. Proteins with an FDR adjusted p-value < 0.05 and a fold change > 2 were considered significant.

### Statistics

GraphPad Prism was used for all statistical analyses described in the text. Quantitative results that required comparisons between groups were subjected to statistical analysis using t-test for two groups or one-way ANOVA for multiple groups, followed by an appropriate ad hoc test. Serum analysis and qualifications were done in a blind fashion wherever possible.

## Acknowledgments

We thank the Cell Biology Histology Core Facility and the Histology and Immunohistochemistry Core of the University of Oklahoma Health Sciences Center (OUHSC) Stephenson Cancer Center for histology and tissue sectioning, supported by P30CA225520 and P20GM103639. We thank the Kansas PKD Research and Translational Core Center (U54DK126126) and the National PKD Research Resource Consortium for normal and ADPKD tissue section slides. We thank IDeA National Resource for Quantitative Proteomics and NIH/NIGMS grant R24GM137786 for quantitative proteomics and analysis. We thank Dr. Christin Vanbeek for helping us with histo-pathology analysis. We also thank Dr. Harini Bagavant for providing us with metabolic cages for characterizing urine parameters. This work was supported by R01DK126705, National Institute of Diabetes and Digestive and Kidney Diseases (NIDDK) and the John S. Gammill Endowed Chair in Polycystic Kidney Disease.

## Author contributions

M.M.P and L.T. designed the study. M.M.P, V.G, and E.P performed the experiments and generated the data. M.M.P and V.G analyzed and interpreted the data. K.Z contributed to data interpretation. M.M.P and L.T wrote the manuscript. All the authors read the manuscript, and their comments and suggestions were incorporated into the manuscript.

## References

1. Bergmann, C. et al. Polycystic kidney disease. Nat Rev Dis Primers 4, 50 (2018).

2. Harris, P.C. & Torres, V.E. Polycystic Kidney Disease, Autosomal Dominant. in GeneReviews((R)) (eds. Adam, M.P. et al.) (Seattle (WA), 1993).

3. Sweeney, W.E. & Avner, E.D. Polycystic Kidney Disease, Autosomal Recessive. in GeneReviews((R)) (eds. Adam, M.P. et al.) (Seattle (WA), 1993).

4. Goggolidou, P. & Richards, T. The genetics of Autosomal Recessive Polycystic Kidney Disease (ARPKD). Biochim Biophys Acta Mol Basis Dis 1868, 166348 (2022).

5. Goksu, S.Y., Leslie, S.W. & Khattar, D. Renal Cystic Disease. in StatPearls (Treasure Island (FL), 2024).

6. Stokman, M., Lilien, M. & Knoers, N. Nephronophthisis-Related Ciliopathies. in GeneReviews((R)) (eds. Adam, M.P. et al.) (Seattle (WA), 1993).

7. Van De Weghe, J.C., Gomez, A. & Doherty, D. The Joubert-Meckel-Nephronophthisis Spectrum of Ciliopathies. Annu Rev Genomics Hum Genet 23, 301–329 (2022).

8. Econimo, L. et al. Autosomal Dominant Tubulointerstitial Kidney Disease: An Emerging Cause of Genetic CKD. Kidney Int Rep 7, 2332–2344 (2022).

9. Wolf, M.T. & Hildebrandt, F. Nephronophthisis. Pediatr Nephrol 26, 181–94 (2011).

10. Wolf, M.T.F., Bonsib, S.M., Larsen, C.P. & Hildebrandt, F. Nephronophthisis: a pathological and genetic perspective. Pediatr Nephrol (2023).

11. Benmerah, A., Briseno-Roa, L., Annereau, J.P. & Saunier, S. Repurposing small molecules for nephronophthisis and related renal ciliopathies. Kidney Int 104, 245–253 (2023).

12. Petzold, F. et al. The genetic landscape and clinical spectrum of nephronophthisis and related ciliopathies. Kidney Int 104, 378–387 (2023).

13. Salomon, R., Saunier, S. & Niaudet, P. Nephronophthisis. Pediatr Nephrol 24, 2333–44 (2009).

14. Shimizu, K., Nihira, N.T., Inuzuka, H. & Wei, W. Physiological functions of FBW7 in cancer and metabolism. Cell Signal 46, 15–22 (2018).

15. Wang, Z. et al. Emerging roles of the FBW7 tumour suppressor in stem cell differentiation. EMBO Rep 13, 36–43 (2011).

16. Davis, R.J., Welcker, M. & Clurman, B.E. Tumor suppression by the Fbw7 ubiquitin ligase: mechanisms and opportunities. Cancer Cell 26, 455–64 (2014).

17. Maskey, D. et al. Cell cycle-dependent ubiquitylation and destruction of NDE1 by CDK5-FBW7 regulates ciliary length. EMBO J 34, 2424–40 (2015).

18. Petsouki, E. et al. FBW7 couples structural integrity with functional output of primary cilia. Commun Biol 4, 1066 (2021).

19. Suryo Rahmanto, A., et al. FBW7 suppression leads to SOX9 stabilization and increased malignancy in medulloblastoma. EMBO J 35, 2192–2212 (2016).

20. Shao, X., Somlo, S. & Igarashi, P. Epithelial-specific Cre/lox recombination in the developing kidney and genitourinary tract. J Am Soc Nephrol 13, 1837–46 (2002).

21. Kang, H.M. et al. Sox9-Positive Progenitor Cells Play a Key Role in Renal Tubule Epithelial Regeneration in Mice. Cell Rep 14, 861–871 (2016).

22. Raza, S. et al. SOX9 is required for kidney fibrosis and activates NAV3 to drive renal myofibroblast function. Sci Signal 14(2021).

23. Kumar, S. et al. Sox9 Activation Highlights a Cellular Pathway of Renal Repair in the Acutely Injured Mammalian Kidney. Cell Rep 12, 1325–38 (2015).

24. Huang, L. et al. TMEM237 is mutated in individuals with a Joubert syndrome related disorder and expands the role of the TMEM family at the ciliary transition zone. Am J Hum Genet 89, 713–30 (2011).

25. Li, D. et al. An Nphp1 knockout mouse model targeting exon 2-20 demonstrates characteristic phenotypes of human nephronophthisis. Hum Mol Genet 31, 232–243 (2021).

26. Jiang, S.T. et al. Targeted disruption of Nphp1 causes male infertility due to defects in the later steps of sperm morphogenesis in mice. Hum Mol Genet 17, 3368–79 (2008).

27. Lancaster, M.A. et al. Impaired Wnt-beta-catenin signaling disrupts adult renal homeostasis and leads to cystic kidney ciliopathy. Nat Med 15, 1046–54 (2009).

28. Attanasio, M. et al. Loss of GLIS2 causes nephronophthisis in humans and mice by increased apoptosis and fibrosis. Nat Genet 39, 1018–24 (2007).

29. Shao, L. et al. Genetic reduction of cilium length by targeting intraflagellar transport 88 protein impedes kidney and liver cyst formation in mouse models of autosomal polycystic kidney disease. Kidney Int 98, 1225–1241 (2020).

30. Gerakopoulos, V., Ngo, P. & Tsiokas, L. Loss of polycystins suppresses deciliation via the activation of the centrosomal integrity pathway. Life Sci Alliance 3(2020).

31. Bai, Y. et al. Primary cilium in kidney development, function and disease. Front Endocrinol (Lausanne) 13, 952055 (2022).

32. Gupta, S., Ozimek-Kulik, J.E. & Phillips, J.K. Nephronophthisis-Pathobiology and Molecular Pathogenesis of a Rare Kidney Genetic Disease. Genes (Basel) 12(2021).

33. Ma, M., Tian, X., Igarashi, P., Pazour, G.J. & Somlo, S. Loss of cilia suppresses cyst growth in genetic models of autosomal dominant polycystic kidney disease. Nat Genet 45, 1004–12 (2013).

34. Hinze, C. et al. Kidney Single-cell Transcriptomes Predict Spatial Corticomedullary Gene Expression and Tissue Osmolality Gradients. J Am Soc Nephrol 32, 291–306 (2021).

35. Mao, J.H. et al. Fbxw7/Cdc4 is a p53-dependent, haploinsufficient tumour suppressor gene. Nature 432, 775–9 (2004).

36. Schulein-Volk, C. et al. Dual regulation of Fbw7 function and oncogenic transformation by Usp28. Cell Rep 9, 1099–109 (2014).

37. Zhang, C. et al. Cyclin-Dependent Kinase 1 Activity Is a Driver of Cyst Growth in Polycystic Kidney Disease. J Am Soc Nephrol 32, 41–51 (2021).

38. Husson, H. et al. Reduction of ciliary length through pharmacologic or genetic inhibition of CDK5 attenuates polycystic kidney disease in a model of nephronophthisis. Hum Mol Genet 25, 2245–2255 (2016).

39. Tao, S. et al. Glycogen synthase kinase-3beta promotes cyst expansion in polycystic kidney disease. Kidney Int 87, 1164–75 (2015).

40. Bukanov, N.O., Smith, L.A., Klinger, K.W., Ledbetter, S.R. & Ibraghimov-Beskrovnaya, O. Long-lasting arrest of murine polycystic kidney disease with CDK inhibitor roscovitine. Nature 444, 949–52 (2006).

41. Watanabe, S. et al. PKD1 mutation may epistatically ameliorate nephronophthisis progression in patients with NPHP1 deletion. Clin Case Rep 7, 336–339 (2019).

42. Sailo, B.L. et al. FBXW7 in Cancer: What Has Been Unraveled Thus Far? Cancers (Basel) 11(2019).

43. Searle, B.C. et al. Chromatogram libraries improve peptide detection and quantification by data independent acquisition mass spectrometry. Nat Commun 9, 5128 (2018).

44. Graw, S. et al. proteiNorm - A User-Friendly Tool for Normalization and Analysis of TMT and Label-Free Protein Quantification. ACS Omega 5, 25625–25633 (2020).

45. Ritchie, M.E. et al. limma powers differential expression analyses for RNA-sequencing and microarray studies. Nucleic Acids Res 43, e47 (2015).

