## Supplemental data for "Nephronophthisis-associated FBW7 mediates cyst-dependent decline of renal function in ADPKD"

Supplemental Figure 1

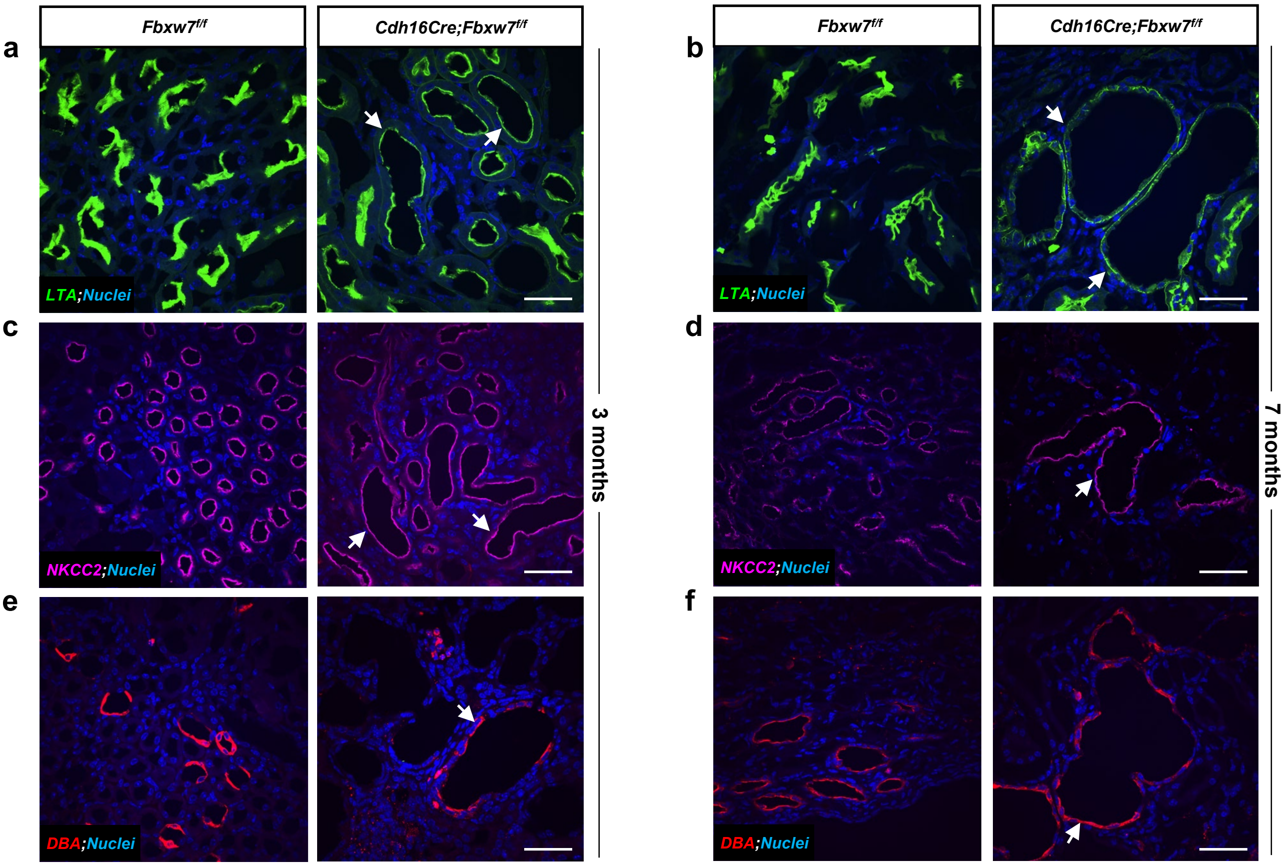

Supplemental Figure 2

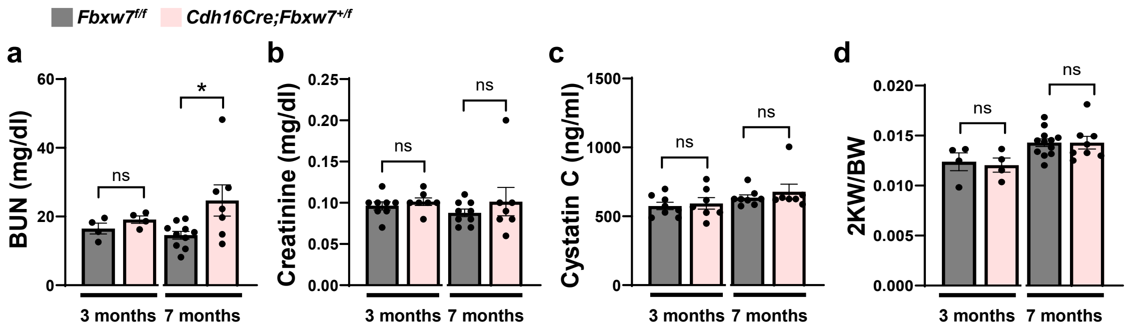

**Supplemental Figure 3**

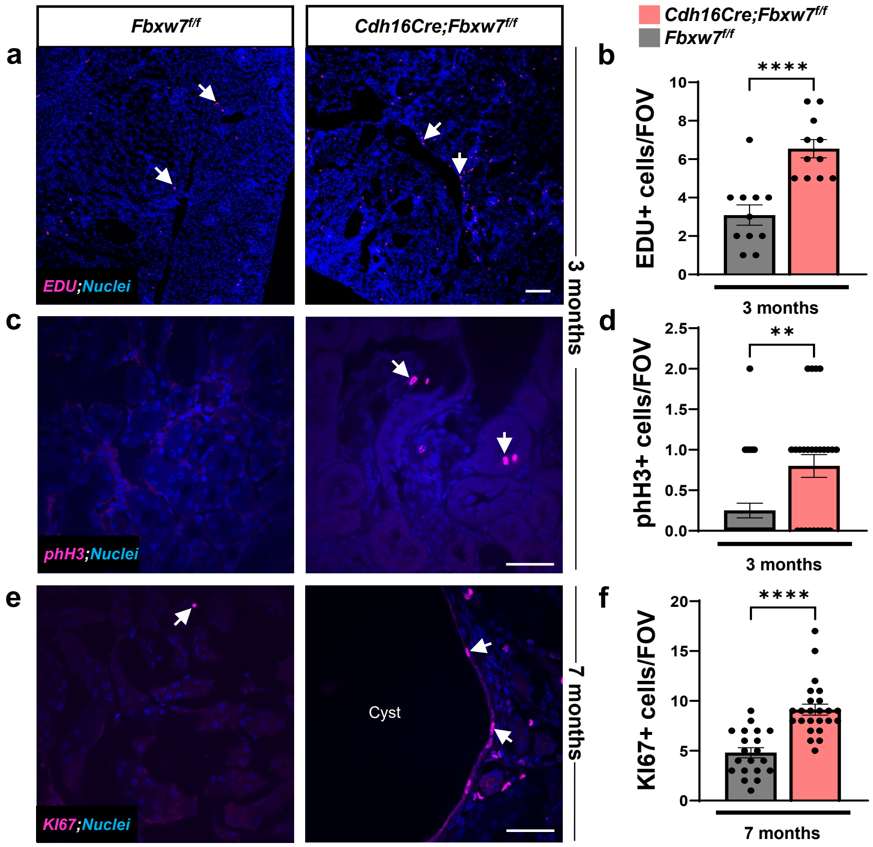

Supplemental Figure 4

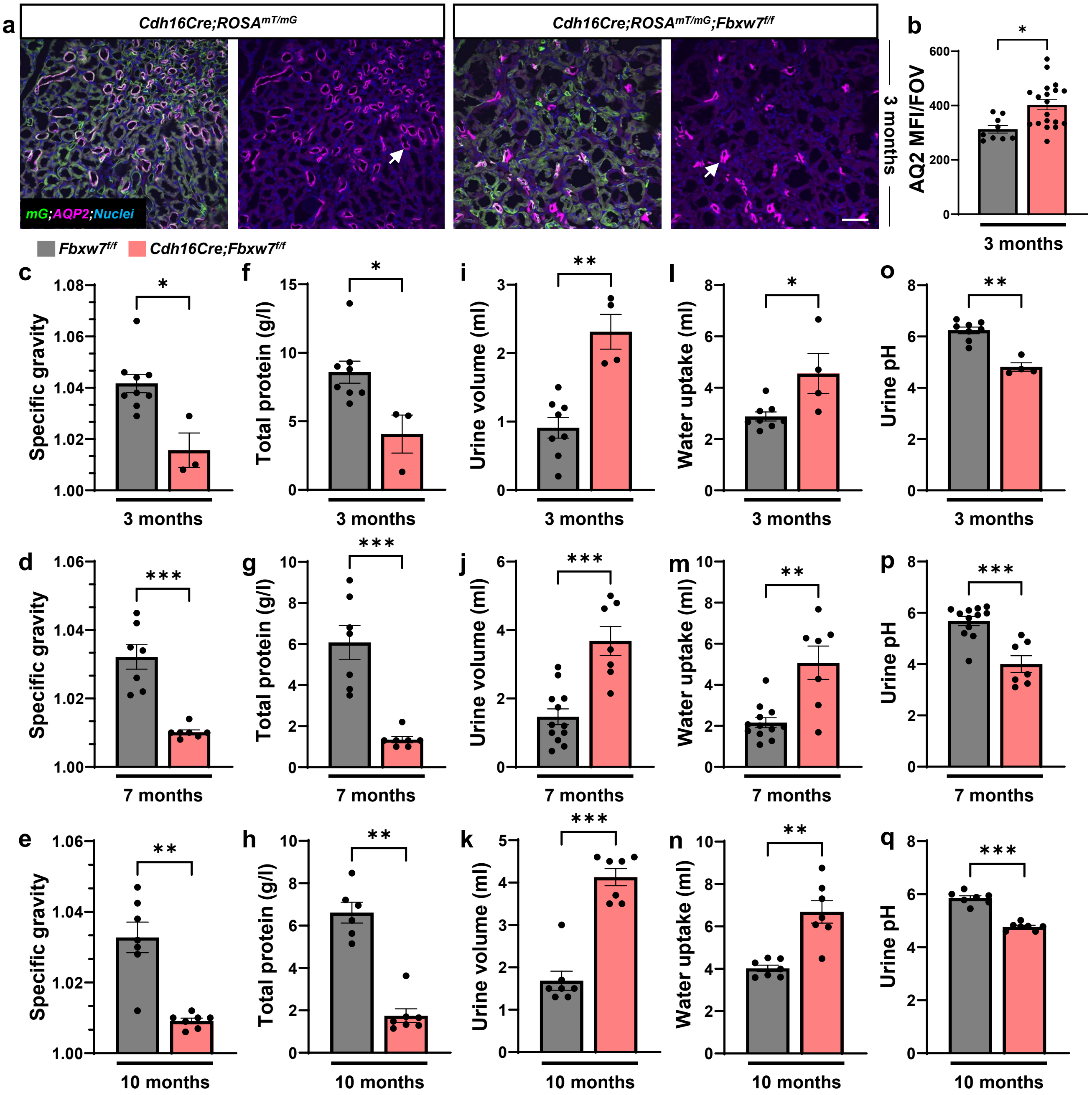

Supplemental Figure 5

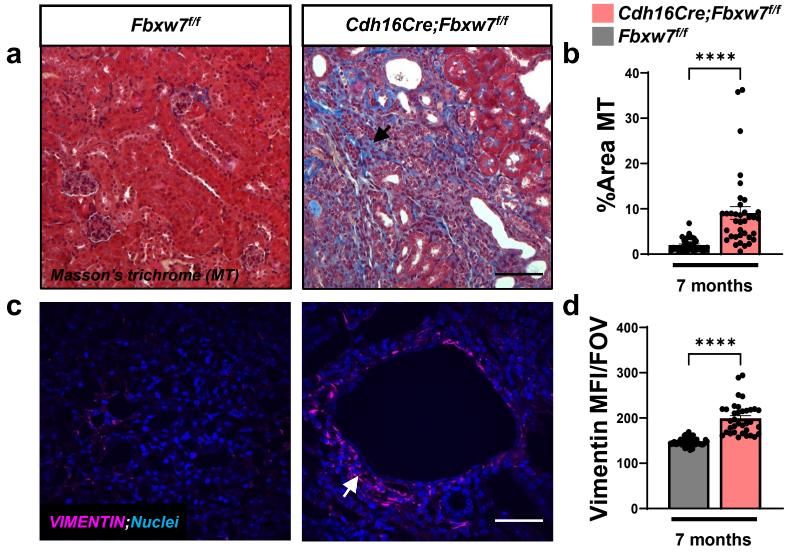

### Supplemental Figure 6

**a** **mIMCD3 Cells: Single cell colony screen for *Fbxw7* null cells**

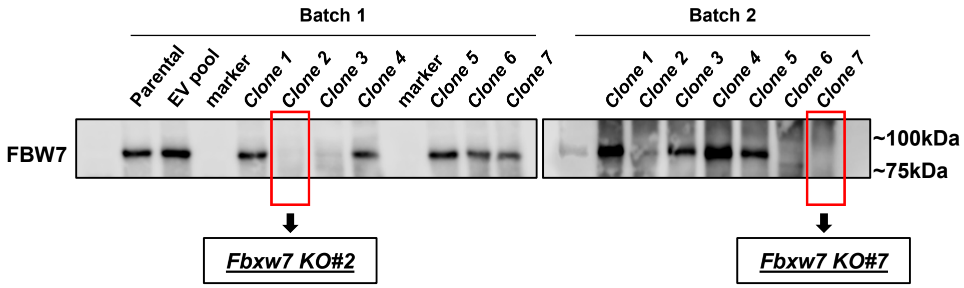

**Supplemental Figure 7**

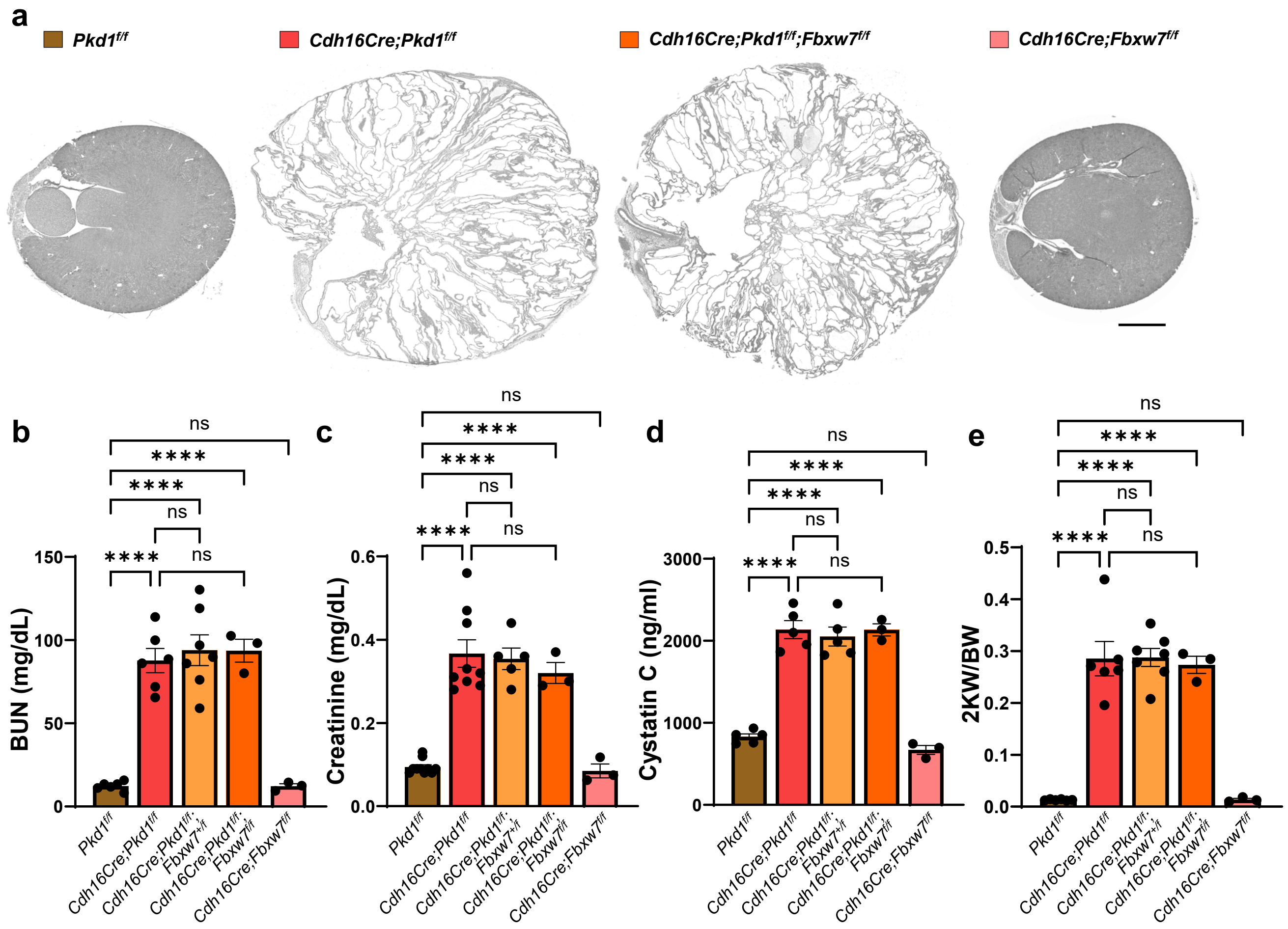

Supplemental Figure 8

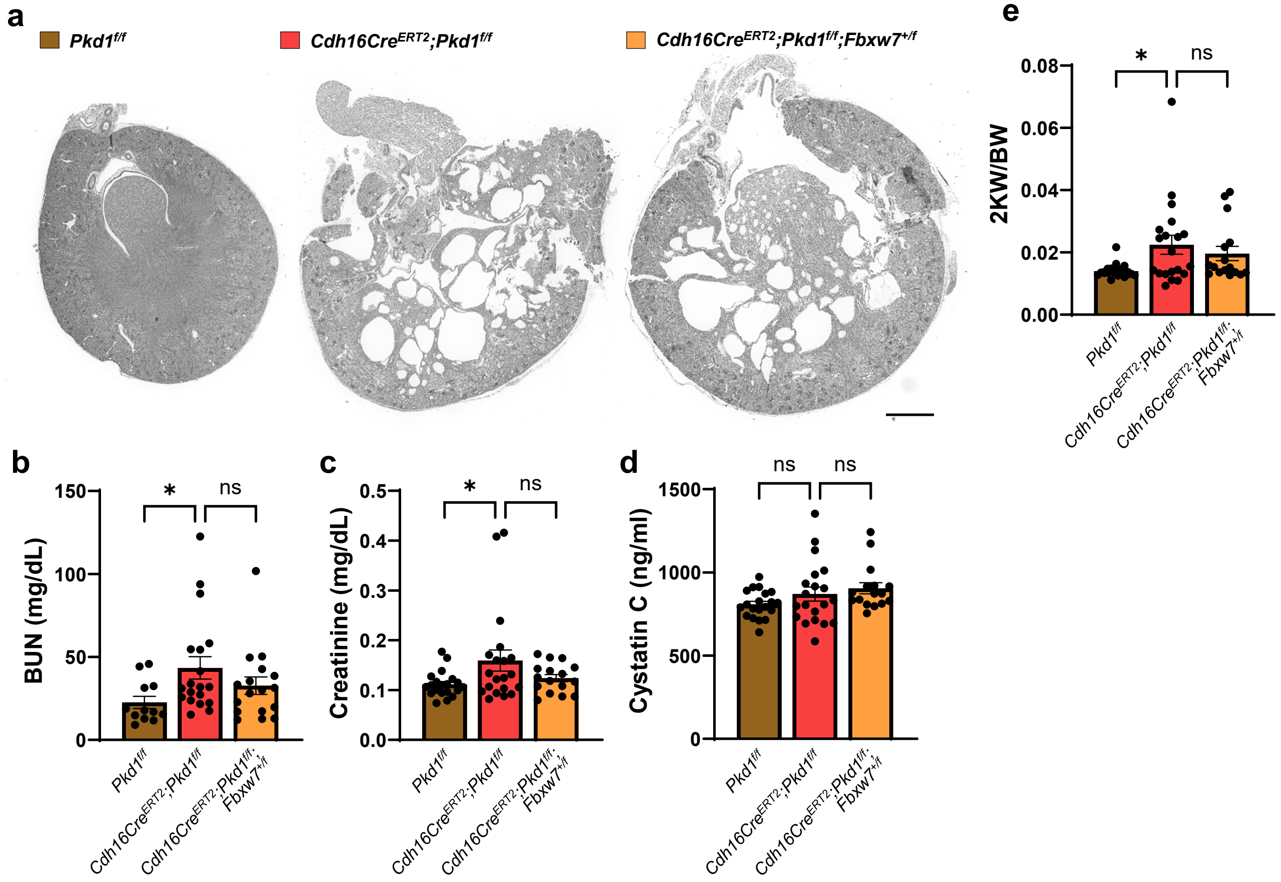

Supplemental Figure 9

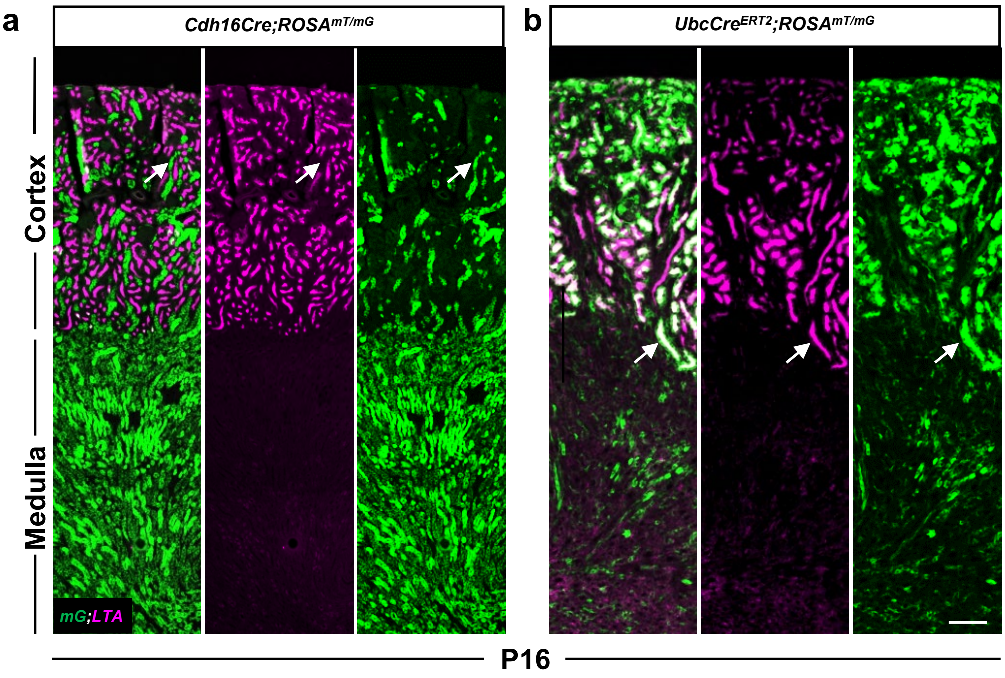

**Supplemental Figure 10**

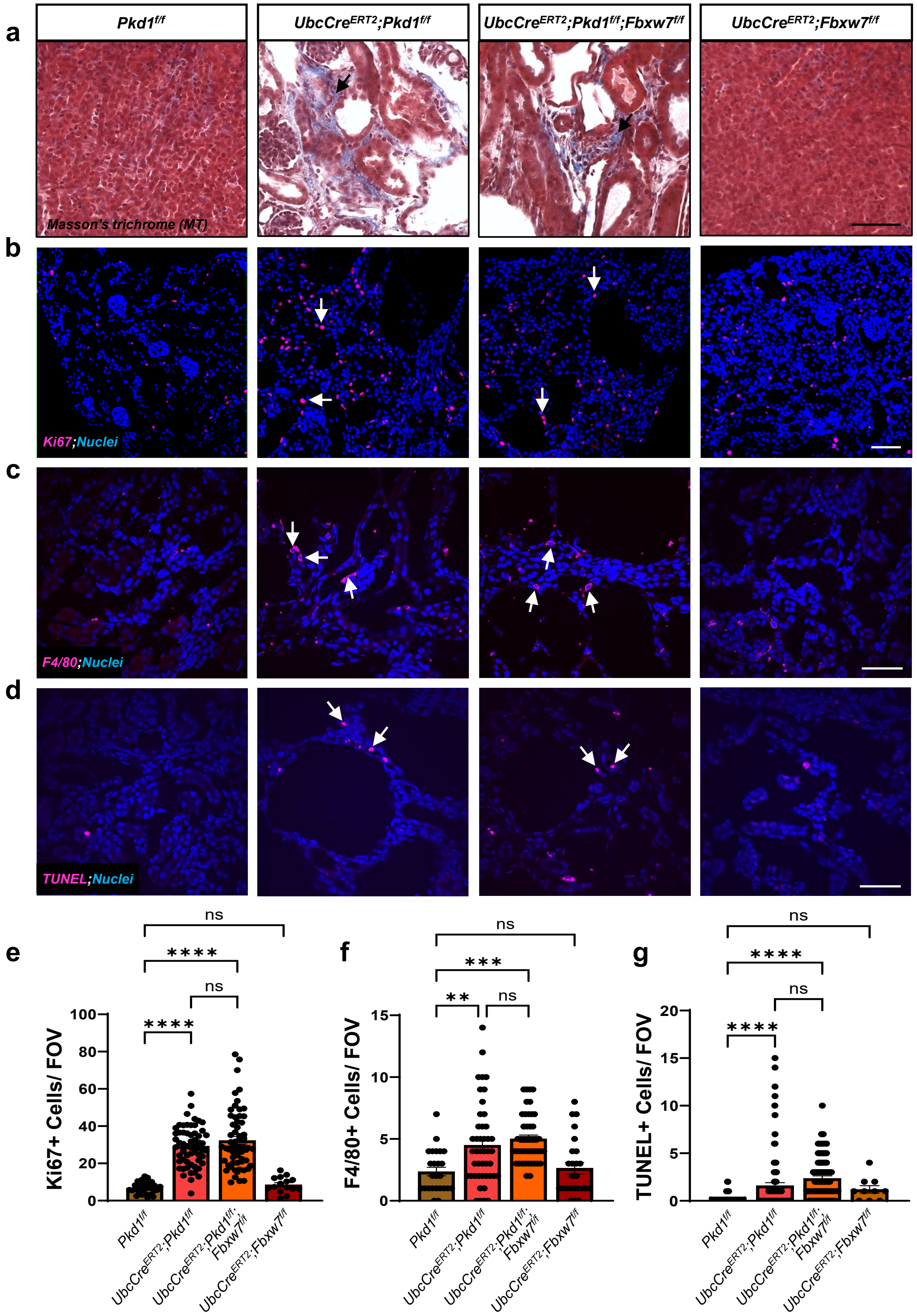

### Supplemental Figure 11

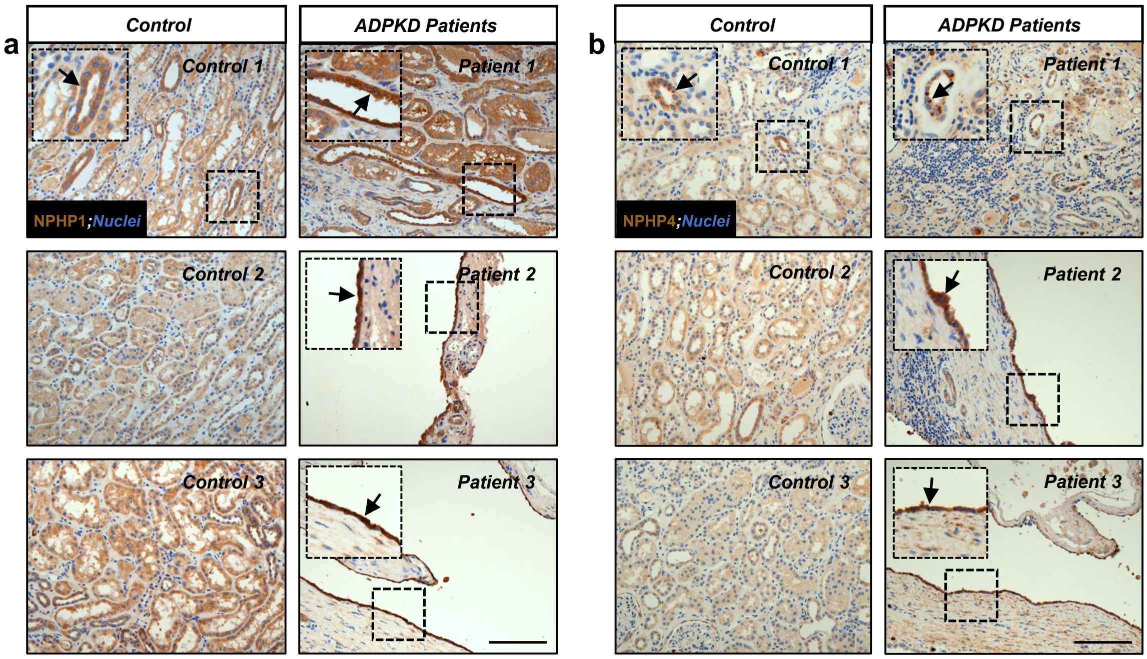

**Supplemental Figure 12**

a

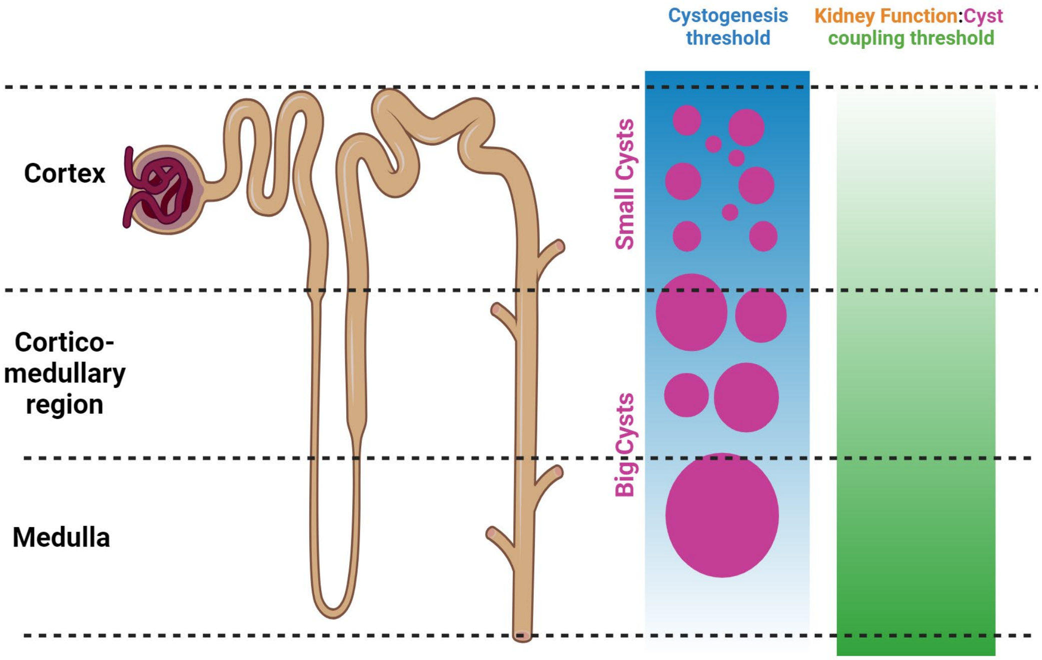

**Supplemental Fig. 1: Deletion of *Fbxw7* results in cyst formation in different tubular segments of the kidney.** Representative images of immunofluorescence staining for (a-b) proximal tubule marker, Lotus tetragonolobus agglutinin (LTA), (c-d) thick ascending limb of the loop of Henle marker, NKCC2, and (e-f) collecting duct marker, Dolichos biflorus agglutinin (DBA) from 3- and 7-month-old kidneys of *Fbxw7*$^{f/f}$ and *Cdh16Cre;Fbxw7*$^{f/f}$ mice. White arrows show dilated or cystic tubular segments of the kidney that are positive for (a-b) LTA (green), (c-d) NKCC2 (pink), and (e-f) DBA (red). Nuclei are stained with DAPI (blue). Scale bar: 50 μm.

**Supplemental Fig. 2: Heterozygous deletion of *Fbxw7* results in mild kidney function decline in aged mice.** (a) Serum BUN, (b) Creatinine, (c) Cytastin C, and (d) 2KW/BW from 3- and 7-month-old *Fbxw7*$^{f/f}$ and *Cdh16Cre;Fbxw7*$^{f/f}$ mice. Each data point represents one animal. All graphs for comparison between *Fbxw7*$^{f/f}$ and *Cdh16Cre;Fbxw7*$^{f/f}$ at different time points are merged to have a common y-axis for comprehensive representation. Statistical analysis was performed using the Mann-Whitney test and is presented as the mean ± SEM (*: $p<0.05$, ns: not significant).

**Supplemental Fig. 3: Deletion of *Fbxw7* increases proliferation in the kidney.** (a-d) Representative images and quantification of (a) EdU (10x magnification) and (c) Phospho-HISTONE H3 (pHH3) immunofluorescence staining from 3-month-old kidneys of *Fbxw7*$^{f/f}$ and *Cdh16Cre;Fbxw7*$^{f/f}$ mice. White arrows show EdU or pHH3-positive (pink) cells. Nuclei are stained with DAPI (blue). Scale bar: (a) 100 μm and (b) 50 μm. (b and d) Each data point represents the number of (b) EdU (60x magnification) or (d) pHH3 positive cells per FOV. For each genotype, n≥10 FOVs were scored from n=3 animals. Statistical analysis was performed using the Mann-Whitney test and is presented as the mean ± SEM (**: $p<0.01$, ****: $p<0.0001$). (e-f) Representative images and quantification of Ki67 immunofluorescence staining from 7-month-old kidneys of *Fbxw7*$^{f/f}$ and *Cdh16Cre;Fbxw7*$^{f/f}$ mice. White arrows show Ki67-positive (pink) cells of the cystic epithelium. Nuclei are stained with DAPI (blue). Scale bar: 50 μm. For each genotype, n≥20 FOVs were scored from n=3 animals. Statistical analysis was performed using the Mann-Whitney test and is presented as the mean ± SEM (****: $p<0.0001$).

**Supplemental Fig. 4: Deletion of *Fbxw7* results in reduced AQUAPORIN2-positive tubules and urine-concentrating defects.** (a-b) Representative images and quantification of immunofluorescence staining for AQUAPORIN2 (AQP2) from 3-month-old kidneys of *Cdh16Cre;ROSA$^{mT/mG}$* and *Cdh16Cre;ROSA$^{mT/mG}$;Fbxw7$^{f/f}$* mice. (a) White arrows show AQP2-positive (pink) tubules where *Cdh16Cre* is active (green). Nuclei are stained with DAPI (blue). Scale bar: 50 µm. (b) Each data point represents MFI per FOV. For each genotype, n≥8 FOVs were scored from n=3 animals. Statistical analysis was performed using the Mann-Whitney test and is presented as the mean ± SEM (*: $p<0.05$). (c-q) Urine-concentrating defects characterized by variations in (c-e) specific gravity, (f-h) total protein concentration in urine, (i-k) urine volume, (i-n)  water uptake, and (o-q) urine pH from 3-, 7-, and 10-month-old *Fbxw7$^{f/f}$* and *Cdh16Cre;Fbxw7$^{f/f}$* mice using metabolic cages. Each data point represents one animal. Statistical analysis was performed using the Mann-Whitney test and is presented as the mean ± SEM (*: $p<0.05$, **: $p<0.01$, ***: $p<0.001$).

**Supplemental Fig. 5: Loss of FBW7 results in excessive tubulointerstitial fibrosis in aged mice.** (a-d) Representative images and quantification of (a-b) MT and (c-d) VIMENTIN staining from 7-month-old kidneys of *Fbxw7$^{f/f}$* and *Cdh16Cre;Fbxw7$^{f/f}$* mice. (a and c) Black and white arrows show MT and VIMENTIN staining, respectively. Scale bar: (a) 200 µm and (c) 50 µm. (b and d) Each data point represents (b) the percent area of MT staining per FOV or (d) MFI per FOV. For each genotype, n≥35 FOVs were scored from n=3 animals. Statistical analysis was performed using the Mann-Whitney test and is presented as the mean ± SEM (****: $p<0.0001$).

**Supplemental Fig. 6: Characterization of *Fbxw7*-null mIMCD3 cell lines.** (a) Immunoblots of FBW7 from different mIMCD3 clones were generated via CRISPR-Cas9 gene editing using

*Fbxw7*-specific sgRNA or EV. Clone 2 from batch 1 and clone 7 from batch 2 were selected and labeled *Fbxw7 KO#2* and *Fbxw7 KO#7*, respectively.

**Supplemental Fig. 7: Deletion of *Fbxw7* in the *Cdh16Cre*-based ADPKD mouse model does not affect kidney function and cystic progression.** (a) Representative images of whole kidney section scan showing cystic progression from *Pkd1*$^{f/f}$, *Cdh16Cre;Pkd1*$^{f/f}$, *Cdh16Cre;Pkd1*$^{f/f}$*;Fbxw7*$^{+/f}$, *Cdh16Cre;Pkd1*$^{f/f}$*;Fbxw7*$^{f/f}$, *Cdh16Cre;Fbxw7*$^{f/f}$ at P16. Scale bar: 400 μm. (b) Serum BUN, (c) Creatinine, (d) Cytastin C, and (e) 2KW/BW from respective genotypes at P16. Each data point represents one animal. Statistical analysis was performed using one-way ANOVA followed by Šídák's multiple comparisons test and is presented as the mean ± SEM (****: p<0.0001, ns: not significant).

**Supplemental Fig. 8: Deletion of *Fbxw7* in the *Cdh16Cre*$^{ERT2}$-based ADPKD mouse model does not affect kidney function and cystic progression.** (a) Representative images of whole kidney section scan showing cystic progression from *Pkd1*$^{f/f}$, *Cdh16Cre*$^{ERT2}$*;Pkd1*$^{f/f}$, and *Cdh16Cre*$^{ERT2}$*;Pkd1*$^{f/f}$*;Fbxw7*$^{+/f}$ at P16. Scale bar: 400 μm. (b) Serum BUN, (c) Creatinine, (d) Cytastin C, and (e) 2KW/BW from respective genotypes at P16. Each data point represents one animal. Statistical analysis was performed using one-way ANOVA followed by Šídák's multiple comparisons test and is presented as the mean ± SEM (*: p<0.05, ns: not significant).

**Supplemental Fig. 9: Expression patterns on *Cre*-mediated recombination in *Cdh16Cre* and *UbcCre*$^{ERT2}$ at P16.** (a-b) Representative images of (a) *Cdh16Cre;ROSA*$^{mT/mG}$ and (b) *UbcCre*$^{ERT2}$*;ROSA*$^{mT/mG}$ kidneys stained with LTA from P16 pups. (a) *Cdh16Cre;ROSA*$^{mT/mG}$ show predominant activity (green) in the medullar region as opposed to the cortex region of the kidney. White arrows show LTA-positive proximal tubules (pink) present in the cortex where *Cdh16Cre* is inactive, and yellow arrows show kidney tubules in the medulla where *Cdh16Cre* is active (green) but are LTA-negative. Orange arrows show a few tubules present in the cortex where *Cdh16Cre* is active but are LTA-negative. (b) *UbcCre*$^{ERT2}$*;ROSA*$^{mT/mG}$ show predominant activity (green) in the cortex region as opposed to the medullar region of the kidney. White arrows show LTA-positive proximal tubules (pink) present in the cortex where *UbcCre*$^{ERT2}$ is active (green), and yellow arrows show a few kidney tubules in the medulla where *UbcCre*$^{ERT2}$ is partially active and LTA-negative. Scale bar: 100 µm.

**Supplemental Fig. 10: Deletion of *Fbxw7* does not ameliorate fibrosis, cell proliferation, inflammation, and apoptosis in ADPKD.** (a) Representative images of MT staining (purple) from *Pkd1*$^{f/f}$, *UbcCre*$^{ERT2}$*;Pkd1*$^{f/f}$, *UbcCre*$^{ERT2}$*;Pkd1*$^{f/f}$*;Fbxw7*$^{f/f}$, and *UbcCre*$^{ERT2}$*;Fbxw7*$^{f/f}$ pups at P16. Black arrows show MT staining (purple). Scale bar: 200 µm. (b-g) Representative images and quantification of immunofluorescence staining for Ki67, F4/80, and TUNEL from respective genotypes at P16. White arrows show (b) Ki67 positive cells, (c) F4/80 positive cells, and (d) TUNEL positive cells (pink). Nuclei are stained with DAPI (blue). Scale bar: 50 µm. (e-g) Each data point represents the number of (e) Ki67-positive, (f) F4/80-positive, and (g) TUNEL-positive cells per FOV. For each genotype, (e) n≥15, (f) n≥25, and (g) n≥10 FOVs were scored from n=3 animals. Statistical analysis was performed using one-way ANOVA followed by Šídák's multiple comparisons test and is presented as the mean ± SEM (**: p<0.01, ***: p<0.001, ****: p<0.0001, ns: not significant).

**Supplemental Fig. 11: ADPKD patients show upregulation of established NPHP genes NPHP1 and NPHP4.** (a-b) Representative images of (a) NPHP1 and (b) NPHP4 expression in normal (n=3) and ADPKD patients (n=3) kidneys using immunohistochemistry. The upper left corner shows a high-magnification image of the insets, and the black arrows show (a) NPHP1 and (b) NPHP4 expression. Scale bar: 1000 µm.

**Supplemental Fig. 12: Diagram showing threshold for coupling cystogenesis to kidney function in different kidney segments.** (a) Tubules in the kidney's cortex are more resistant to cyst formation than those in the medulla, yet they have a greater sensitivity in linking cyst development to the loss of kidney function.
